# Poaching exacerbates the effects of climate change on the long-term viability of an endemic South African succulent plant species

**DOI:** 10.1101/2025.01.20.633962

**Authors:** Arjan Engelen, Katrina Davis, Allan G. Ellis, Roberto Salguero-Gómez

## Abstract

1. Natural populations often face multiple threats that jeopardise their viability, yet most population forecasts consider only single threats, such as shifts in precipitation or temperature. To better understand ecological and conservation challenges, incorporating multiple threats is essential. South African endemic dwarf succulents, for example, are under dual threats from climate change and illegal harvesting for the ornamental plant trade.
2. We developed a stochastic integral projection model (IPM) using demographic data from 1999-2003, for 4776 individuals of the dwarf succulent, *Argyroderma pearsonii*, to evaluate the impacts of climate change and harvesting on its short-term (<40 years) extinction risk and long-term viability. We built simulations to explore 90 scenarios combining climate conditions (historical, mild climate change [SSP126], and moderate climate change [SSP245]), harvesting methods (fruit harvesting, non-selective plant harvesting, or size-selective plant harvesting), and harvest frequencies (annual to once every decade). We supplemented our approach with perturbation analyses to identify critical demographic components influencing population persistence.
3. We show that historically *A. pearsonii* populations have been stable or growing. Populations are predicted to remain viable under mild climate change, but moderate climate change results in declines in approximately 50% of simulated populations, though extinctions are not projected within 40 years without harvesting. In contrast, plant harvesting significantly increases extinction risk, especially size-selective harvesting of mature individuals, which disrupts demographic components critical to population growth. Even harvesting as infrequently as once per decade under historical climates leads to declines, with climate change exacerbating these effects.
4. **Synthesis**: Illegal harvesting poses a critical threat to biodiversity, as shown in this case study of a South African endemic succulent. Our findings indicate that plant harvesting is unsustainable under any scenario, particularly with projected climatic pressures. However, fruit collection emerges as a potentially sustainable alternative to meet the ornamental plant trade’s demands. Our study highlights the importance of incorporating multiple threats into population forecasts, providing valuable tools for ecologists and conservation managers.

## Introduction

Humans are profoundly transforming the planet (Venter *et al*., 2016) and negatively impacting biodiversity (Bellard *et al*., 2014). Our activities increase extinction risk through climate change-induced losses of suitable niche space (Musil *et al*., 2005; Young *et al*., 2016; Manes *et al*., 2021), and direct impacts through land use change (Davison *et al*., 2021), invasive alien species (Bellard *et al*., 2016), and overexploitation (Hinsley *et al*., 2023). Moreover, we are becoming increasingly aware of compounding impacts from multiple stressors (Co□té *et al*., 2016), suggesting that consideration of multiple threats is essential for estimating resilience to anthropogenic change.

Desert plant species may be especially vulnerable to multiple anthropogenic threats (Pillet *et al*. 2022). Extreme climatic conditions in deserts often hold plant populations near their thresholds of physiological tolerance (Musil *et al*., 2005, 2009; Grey and Atkinson, 2024), while edaphic specialisation and dispersal lags may render populations incapable of tracking suitable climatic envelopes under climate change (Foden *et al*., 2007; Young *et al*., 2016; Eibes *et al*., 2022). Furthermore, desert plants are popular items of trade in legal and illegal horticultural markets (Margulies *et al*., 2020), and overharvesting of wild populations has become an important conservation concern for arid-adapted plant taxa (Goettsch *et al*., 2015; Young *et al*., 2016; Frantz *et al*., 2021).

While wild-harvested desert plants from South Africa have been traded internationally since the 1970s (Jarvis, 1979), demand for dwarf leaf succulent species, particularly Aizoaceae from the Succulent Karoo, has increased considerably since 2019 (*cf* Hammer, 1997; Frantz *et al*., 2021). Limited conservation infrastructure and economic opportunities in the Succulent Karoo is thought to have incentivised the rise in illegal harvesting in recent years (Frantz *et al*., 2021; Acker-Cooper *et al*., 2024), with severe impacts for several endemic species (SANBI, 2024). Coupled with ongoing climate change, continued exploitation of these plants for the horticultural trade may compromise their resilience to anthropogenic stressors. Therefore, appropriate conservation targets are needed to ensure sustainable harvesting practices under climate change (Frantz *et al*., 2021).

Past studies on anthropogenic impacts to biodiversity have primarily focused on climatic stressors using distribution-modelling approaches (Sinclair *et al*., 2010). While these methods are valuable, they often fail to account for the combined effects of climate change and direct human pressures, which may interact synergistically to amplify impacts (Bellard, 2012; Bellard *et al*., 2014; Mantyka-Pringle *et al*., 2015). Furthermore, distribution-modelling approaches have been criticised for their inability to establish clear links between specific ecological processes and population viability (Briscoe *et al*., 2019). To address this gap, process-based evidence is needed to better understand the cumulative effects of multiple anthropogenic stressors, such as climate change and wild harvesting, on populations (Bellard *et al*., 2012). Adopting a multifaceted approach that integrates ecological, socio-economic, and environmental contexts could lead to more effective population management strategies (Davis, 2022; Hubschle and Margulies, 2024).

Structured population models are flexible tools for projecting the impacts of multiple anthropogenic stressors on biodiversity (Merow *et al*., 2014; Levin *et al*., 2021). These models track changes in population size and structure as a function of how one or more continuous (*e.g.*, size) and/or discrete variables (*e.g.*, age) predict the vital rates of survival, development, reproduction, and recruitment (Easterling *et al*., 2000; Caswell *et al*., 2001). When vital rates have been assessed over multiple time steps, structured population models can be parameterised to forecast population dynamics along a stochastic trajectory of environmental change, derived from field observations or global climate models, while examining the impacts of multiple threats (Keith *et al*., 2008; Williams *et al*., 2015; Davis, 2022). Additionally, the vital rates of these models can be ‘perturbed’ to explore population-level sensitivities to additional impacts, such as culling (Bogdan *et al*., 2021; Davis, 2022), or fruit harvesting (Freckleton *et al*., 2003; Venter and Witkowski, 2013), making them ideal to explore ‘what if’ scenarios.

Here, we parameterise an Integral Projection Model (IPM; Easterling, 2000) to project the demographic responses of the South African dwarf succulent *Argyroderma pearsonii* (Aizoaceae) to the combined pressures of climate change and plant/fruit-capsule (*i.e.*, seed) harvesting. To do so, we use five years of data collected across six sites in the Knersvlakte region of the Western Cape of South Africa (Figure 1A, B). Specifically, we hypothesise that: (H1) Climate change will reduce population growth rates and increase extinction risks compared to historical baselines, as previous studies show higher mortality in *Argyroderma* species under warming (Musil *et al*., 2005; 2009) and drought (Grey and Atkinson, 2024). (H2) Plant harvesting and fruit capsule harvesting will decrease the population growth rate and increase extinction risk with increasing frequencies of harvesting events. (H3) Plant harvesting will more strongly affect population growth than harvesting limited to fruit capsules. *A. pearsonii* is iteroparous and produces thousands of seeds per fruit capsule (Hartmann, 2004; Ellis *et al*., 2007). Therefore, while fruit harvesting may reduce the fraction of new recruits from seed, it is not expected to impact future reproduction, nor is it expected to severely impact the regenerative capacities of populations under favourable conditions. Furthermore, (H4) plant harvesting will have a stronger effect on the population growth rate when harvesters target larger plants than when plants are harvested non-selectively because larger individuals are the primary source of new recruits (Ellis *et al*., 2007). Finally, we hypothesise (H5) that future climate change will exacerbate the impacts of increasing harvest frequencies on the population growth rate.

**Figure 1.**
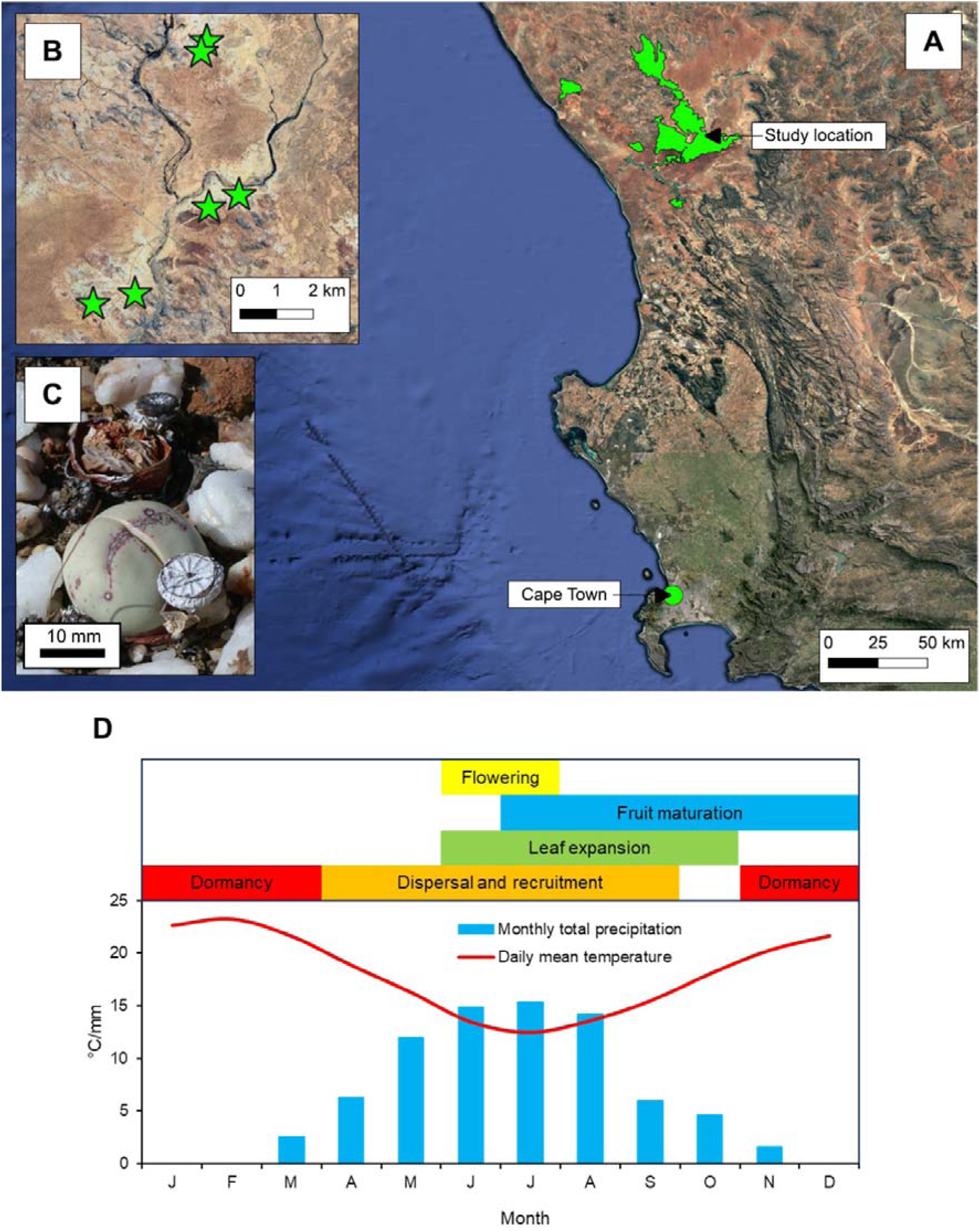
Study sites and phenology of our study species, *Argyroderma pearsonii*, used to examine the effects of harvesting and climate change on the species’ population viability. (A) Study location in the Knersvlakte Quartz Vygieveld of South Africa (31.375□S, 18.550□E). Delimitation of the Knersvlakte Quartz Vygieveld was taken from the National Vegetation Map of South Africa (2012). (B) Six sites were monitored annually during October-November of 1999 to 2003. (C) *A. pearsonii*, the focal plant species of this study. (D) Annual climate of the Knersvlakte quartz fields and phenology of *A. pearsonii*. Monthly total precipitation and daily mean temperature data were digitised from Mucina *et al*. (2006).

## Methods

### Study system and species

Our study was conducted in the Knersvlakte Quartz Vygieveld (∼31.4^□^S, 18.7^□^E, ∼250 m.a.s.l.), part of the hyperdiverse Succulent Karoo biome of South Africa (Mucina *et al*., 2006). The region experiences a mean annual temperature of 18.1°C, with mean daily maxima ranging from 19.2-31.5°C throughout the year (Mucina *et al*., 2006). Total annual precipitation is 116 mm, with 84% of precipitation occurring in the autumn and winter months from March to August (Mucina *et al*., 2006). Over the past century, the frequency of hot days has increased, without discernible changes to annual rainfall patterns (Davis *et al*., 2016). Future projections indicate rising temperatures (Hulme *et al*., 2001), with drying induced by the southward migration of mid-latitude frontal systems (MacKellar *et al*., 2007).

Our study species, *Argyroderma pearsonii* (N.E.Br.) Schw (Aizoaceae) (Figure 1C), is an endemic perennial dwarf succulent, usually comprised of a single pair of rounded, appressed leaves (Figure 1C, Hartmann, 2004). Rarely, individuals may branch to form two (0.68% in our dataset), three (0.14%), or even four (0.08%) leaf pairs (Hartmann, 2004). The phenology of *A. pearsonii* is aligned with the winter rainfall cycle of the Knersvlakte (Fig 1D). Dormancy occurs during the dry season (November-April; Hartmann, 2004), while winter rains in April stimulate new leaf and flower development. Mature individuals bloom from mid-May to mid-June (Ellis *et al*., 2006). Typically, one solitary hermaphroditic flower emerges from the terminus of the leaf pair per season (Hartmann, 2004), though our dataset indicates that individuals with multiple leaf pairs may produce additional flowers. *A. pearsonii* only reproduces sexually, is self-incompatible, and relies on insect-pollination (Hartmann, 1977; Ellis *et al*., 2007; Boucher *et al*., 2024). Fruit maturation and leaf growth occur in parallel from June to November, with fruit maturation extending into December. Seeds, housed in hygrochastic fruit capsules, are dispersed during heavy rainfall the following winter (Parolin, 2006; Schmiedel *et al*., 2021). Seedling germination and growth proceeds throughout the rainy season (April-September) (Figure 1D).

### Demographic data collection

To investigate the combined effects of climate change and wild harvesting on *A. pearsonii*, we used morphometric and life history records for six populations described elsewhere (Ellis and Weis, 2006; Ellis *et al*. 2006; Ellis *et al*., 2007). These populations were distributed in pairs across three drainage basins (Figure 1B) and tracked over five annual censuses from 1999 to 2003. Censuses were conducted at the end of the growth period (October-November) in three to four 1×1 m^2^ plots per site. Plot maps were created to track individuals across years, with an average of 376 ± 262 (S.D.) individuals recorded per site annually (Table S1).

Data collected included individual survival, leaf sizes, branching events, flowering and fruit-set, seed production, and recruitment. Survival and recruitment were determined by systematic searching of the mapped sampling plots. Individuals were scored as alive when they had produced at least one new leaf pair during the growth season. Death was recorded when individuals had a shrivelled, brown leaf pair, or when the plants were killed by herbivory after producing a new leaf pair. Established recruits were defined as individuals first detected after 1999, possessing a single turgid leaf pair. Developmental states were described using total leaf size. For individuals with a single leaf pair, this was the maximum diameter across the exposed portion of the new leaf pair (Figure 2). For individuals with multiple leaf pairs, total leaf size was the sum of the diameters of all leaf pairs.

**Figure 2.**
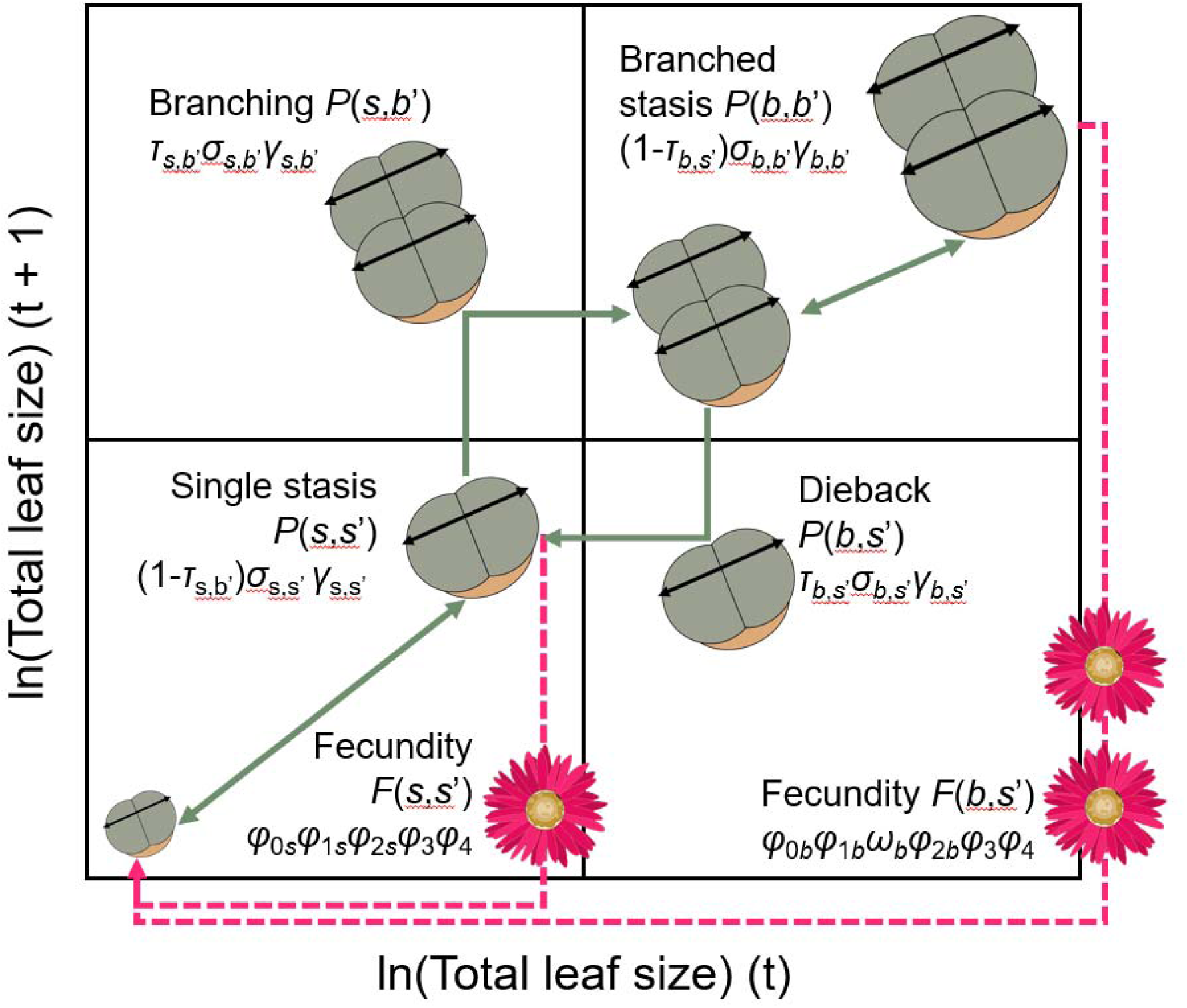
Conceptualisation of the Integral Projection Model (IPM) used to simulate the demographic responses of *Argyroderma pearsonii* to harvesting pressure and projected climate change. This model comprises a two-stage structure for individuals with a *s*ingle leaf-pair (state *s*) and those which *b*ranched to form multiple leaf pairs (state *b*). The model is comprised of four *P*-kernels (rectangular panels), parameterised by survival (σ), growth (γ) and transition probabilities (τ) within and between states *s* and *b* from time *t* to time *t* + 1 (solid arrows). In addition, the model incorporates two *F*-kernels describing contributions of single and branched individuals in time *t* to the recruitment of single-pair individuals in *t*+1 (dashed arrows). The F-kernels were parameterised by functions for flowering (*φ*_0_), fruit set (*φ*_1_), fruit quantity in branched individuals (ω), total seed number (*φ*_2_), recruitment from seed (*φ*_3_) and recruit sizes (*φ*_4_). One variable, ln(Total leaf size), was used to describe the state of individuals in both stages, and is depicted as the double-headed arrows drawn across the leaf pairs in the plant diagrams.

Reproductive success was quantified from the incidence of flowering, fruit set, and seed production. Individuals were scored as having flowered and set fruit if they produced at least one new fruit capsule. Those retaining the withered remains of a flower, but lacking fruit capsules, were scored as having flowered only. Seed production was estimated using allometric equations predicting seed count per fruit capsule based on the maximum diameter of the capsule (see Supplementary Material S2 for details). Allometric predictions and subsequent statistical analyses were conducted in R version 4.2.3 (R Core Team, 2023).

### Harvesting pressure

To explore the effects of plant and fruit harvesting on the population growth rate of *A. pearsonii*, we adjusted records in the original demographic dataset of size-dependent and size-independent survival/mortality, and size-independent fruit set between seasons. This created three modified datasets representing two plant harvesting scenarios and one fruit harvesting scenario (Figure S3), which would be implemented in our population simulations whenever populations experienced a harvesting event.

For the size-dependent plant harvesting scenario (modified dataset 1), we reduced the survival probability between seasons by 90% for individuals with leaf sizes >10 mm. This threshold was chosen because illegal harvesters target mature plants, which are easier to transport, replant, and sell (Margulies, 2020). Mature *A. pearsonii* individuals (>10 mm) are more likely to flower and survive than smaller individuals (Figure S3) and are also more visible in their natural habitat (A.G. Ellis. pers. obs.). We assumed harvesters would maximise yield once a population had been located by targeting 90% of mature individuals (*i.e.*, 1,303 individuals in the original dataset).

A size-independent plant harvesting scenario (modified dataset 2) served as a baseline to assess the effects of size-dependent harvesting on the demographic profile. Here, the same number of individuals (1,303) were removed from the population without regard to size. For both plant harvesting scenarios, harvested individuals were assumed to contribute no reproduction in the year of harvesting, with their flowering scores set to zero.

For the fruit harvesting scenario (modified dataset 3), only fruit set scores were modified, while survival and flowering scores remained unchanged. Assuming harvesters were non-selective regarding leaf or capsule size, the probability of fruit set was reduced by 90% across all sizes. This resulted in the removal of 379 fruit capsules per harvesting event.

### Historical and projected climate variables

We characterised the climatic conditions associated with our demographic dataset to simulate population dynamics under climate change (Davis, 2022). Hindcasted (1950–2022) and projected (2016–2098) monthly mean air temperatures (2 m above ground level) and total precipitation data were obtained from the Copernicus Climate Data Store (2021). Hindcasted data came from ERA5-Land reanalysis (Muñoz-Sabater *et al*., 2021) using the *download_ERA* function in the ‘KrigR’ R package (Kusch & Davy, 2022). Projected data were sourced from the CMIP6 UKESM1-0-LL global climate model (Mulcahy *et al*., 2022) under two shared socio-economic pathways: SSP126 (mild, +2.6°C by 2100) and SSP245 (moderate, +4.5°C by 2100). The UKESM1-0-LL model was chosen for its alignment with conditions experienced by *A. pearsonii* during the census years, while still projecting climatically extreme years. The hindcasted and projected climate datasets exhibited two continuity issues, which were adjusted before proceeding with further analyses (details in the Supplementary Material S4).

To link continuously varying climatic data to a set of discrete annual population states, we classified years along axes of mean annual temperature (MAT) and total annual precipitation (TAP) for the hindcasted, SSP126, and SSP245 datasets. Years were categorised as hot-dry, hot-wet, cool-dry, or cool-wet, with thresholds centred on median MAT and TAP values from the census years (Figs. S4.1, S4.2). This centring ensured that each possible set of climatic conditions was represented by one demographic transition (*i.e.*, a sequential pair of census years) during the population simulations.

### Demographic modelling

To test our hypotheses, we parameterised an Integral Projection Model (IPM, Easterling *et al*. 2000) using demographic data for *A. pearsonii*. IPMs describe population dynamics in discrete time steps using kernel density functions based on a species’ vital rates. Vital rates represent individual survival (σ), growth (γ), and reproduction (*φ*) in time t+1, as functions of a continuous state variable *z* (total leaf size) at time *t* (years). The standard IPM consists of two kernels: (1) the *P* kernel, describing the probability of individuals in state *z* at *t* surviving and transitioning to state *z*’ at time *t*+1, and (2) the *F* kernel, describing the state distribution of new recruits *z*’ at *t*+1 based on reproductive outputs of individuals in state *z* at *t*. For models with multiple life stages (Figure 2), additional kernels handle transitions between discrete states (*e.g.*, from a single leaf pair to multiple leaf pairs; Zambrano & Salguero-Gómez 2014).

Our goals with this IPM were to estimate the stochastic population growth rate (λ*_s_*) and population sizes (*n*) of *A. pearsonii* under hindcasted climates, and mild and moderate projected climate change (H1 and H3), as well as under size-dependent, and size-independent, plant harvesting, and fruit harvesting (H2, H4, and H5). To these ends, our IPM was constructed from vital rate models that explicitly considered variance in all vital rates between years, and between our three harvesting scenarios. Census year (*t*) was a fixed effect in all vital rate mean-models, enabling simulations of stochastic dynamics under the three climate scenarios, while harvesting scenario (θ) was treated as a fixed factor in the survival and fruit set regressions. Site was treated as a random factor in all mean models to account for spatial variation in vital rates. Variance models for size distributions (see ε_γ_ and ε_*φ*_*_4_* below) excluded year and site-effects.

We used generalised linear or additive mixed effects models to describe the survival (*σ*), mean growth (*μ*_γ_), growth variance (*ε*_γ_), flowering probability (*φ_0_*), fruit set probability (*φ_1_*) and seed production (*φ_2_*) at *t*+1 as functions of *z* = ln(total leaf size) at *t* (Fig 2). Recruit establishment (*φ_3_* = recruits/total seeds) and new recruit size distributions (*μ*_*φ*_*_4_, ε*_*φ*_*_4_*) in *t*+1 modelled population regeneration between years, but were not linked to parent sizes due to feasibility constraints. The vital rate regressions were performed using the R packages ‘glmmTMB’ (Brooks *et al*., 2017), ‘mgcv’ (Wood, 2011), ‘gamlss’ (Rigby and Stasinopoulos, 2005), and ‘gamlss.tr’ (Stasinopoulos and Rigby, 2024). Further parameterisation of the vital rate models was informed by an intensive AIC- and pseudo-*R*^2^-based model selection procedure (Supplementary Material S5).

Model selection revealed vital rate differences between single-leaf-pair and branched individuals. To capture these differences, our IPM contained two continuous state classes: *z = s* (*s*ingle leaf pair) and *z = b* (branched, multiple leaf pairs). Transitions (τ) within and between states *s* and *b* were modelled according to the probabilities of: (1) retaining a single leaf pair across times *t* and *t*+1 (*s, s’*); (2) branching (*s, b’*); (3) retaining multiple leaf pairs (*b, b’*); and (4) reverting to a single leaf pair (*b, s’*). As for the vital rate models, the models for τ depended on ln(total leaf size), year (fixed intercepts), and site (random intercepts). The transition models were subjected to the same model selection procedure as the vital rate models (Supplementary Material S5).

We synthesised vital rate models into an IPM using the ‘ipmr’ package (Levin *et al*. 2021). The IPM consisted of a series of mega-kernels (*M*(*z,z’*)), each describing the survival, growth, discrete state transition probabilities, and reproduction of *A. pearsonii* in one of the four demographic time-steps (*t*) recorded during the census period. Each mega-kernel was formed from the concatenation of four kernels (*K*). Each *K* kernel encompassed one of the possible transitions from states *z* = *s* and *z* = *b* in time *t*, given harvesting scenario θ, to states *z*’ = *s*’ and *z*’ = *b*’ in *t*+1, where *t □* [1999; 2002]:

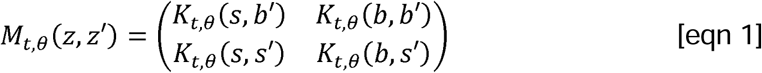

In turn, each *K* kernel was comprised of one *P* sub-kernel, describing the states of individuals between time-steps as the product of the survival, growth, and discrete transition probabilities parameterised by the vital rate regressions for each harvesting scenario. Additionally, the kernels capturing individuals that retained a single leaf pair between time-steps *K*(*s,s’*) and those that transitioned from multiple leaf pairs to a single leaf pair *K*(*b,s’*) each contained an *F* sub-kernel describing the recruitment and size distribution of single leaf pair individuals from the respective reproductive outputs of single and branched individuals. As such, the *K* kernels were defined as follows:

For individuals that remained in stasis as a single leaf pair or were recruited from the seed of single pair individuals from time *t* to *t*+1:

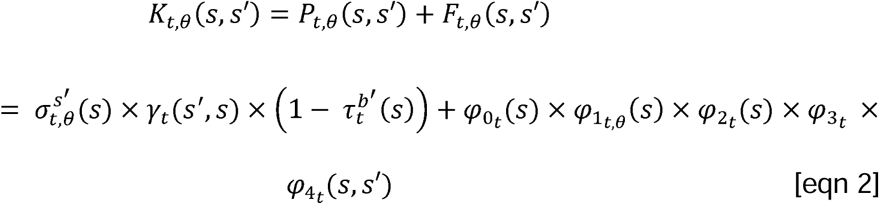

For individuals that branched to have multiple leaf pairs from *t* to *t*+1:

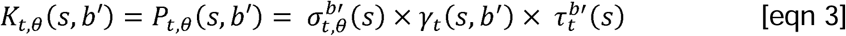

For individuals remaining in stasis as multiple leaf pairs between *t* and *t*+1:

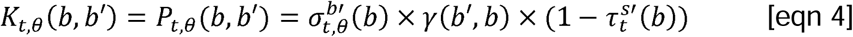

For individuals that reverted to a single leaf pair or were recruited from the seed of branched individuals from *t* to *t*+1:

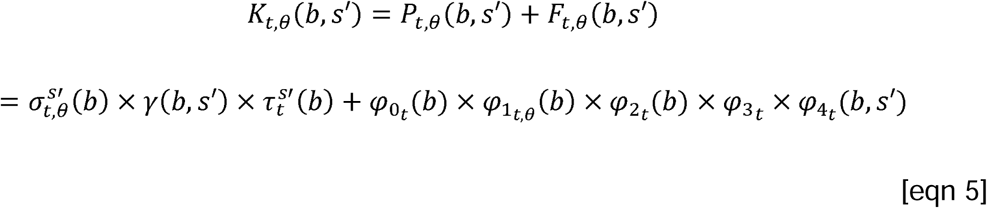

With this model structure, the number of individuals *n* of size *z*’ in time *t+*1 was given by:

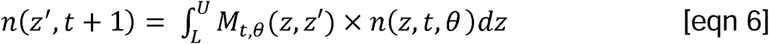

where *L* and *U* represent the respective minimum and maximum values of ln(total leaf size) for states *z* = *s* and *z* = *b* within their relevant sub-kernels (Easterling *et al*., 2000). To avoid the accidental eviction of individuals from the observed range of ln(total leaf size) during the integration, the functions describing the Gaussian probability density of *z*’ in *t*+1 (*i.e.*, the γ and *φ*_4_ functions in equations 2-5) were truncated at *L* and *U* for the relevant state (*s’* or *b’*) in each sub-kernel (Ellner *et al*., 2016). Following integration, each continuous *K*-kernel was discretised using the midpoint rule (Easterling *et al*. 2000) across 100 mesh points per state.

### Population simulations

We conducted stochastic population simulations with our IPM to assess the impacts of climate change and harvesting on the long-term population growth rate λ*_s_*, and short-term extinction risk of *A. pearsonii*. Simulations were performed under three climate scenarios (historical, SSP126, SSP245), and three harvesting scenarios (size-selective plant harvesting, non-selective plant harvesting, and fruit harvesting). To evaluate the maximum sustainable frequencies of harvesting events, we varied the probability of harvesting events from 0 to 1 in intervals of 0.1 for each scenario, resulting in 90 unique combinations of climate, harvest type and frequency. Each scenario was subjected to 10 replicate simulations.

For each simulation, we generated a sequence of 100 kernels (*i.e.*, annual demographic transitions) through which we projected an initial population vector. Kernel sequences were generated using probabilistic sampling of the set of 16 kernels comprising each climate-classified demographic transition (*i.e.*, kernels for warm-dry, cool-dry, warm-wet, and cool-wet years) and harvesting event type (no harvest, plants size-selective, plants non-selective, and fruit), with sample weights depending on the harvesting and climate scenario in question. The initial population vector corresponded to the probability density functions of ln(total leaf size) averaged across all sites and years, multiplied by a randomly sampled plant density value (individuals/m^2^) taken from the census plots.

To implement each harvesting scenario in our kernel sequences, harvesting events were assigned to kernel positions using binomial sampling (harvest/no harvest) weighted by harvesting frequency. Kernels constructed from the appropriate modified dataset were used at positions in the sequences that were assigned for harvest, while kernels constructed from the unmodified dataset were used at positions assigned for no harvest.

To implement each climate scenario in our kernel sequences, annual climate classifications were assigned to kernel positions using three Markovian transition matrices—one for each climate dataset (hindcasted, SSP126, SSP245). The Markovian matrices assigned annual climatic states to kernel sequences according to the probabilities of transitioning to one of the four possible climate states *y* (*e.g.*, warm-dry, cool-dry, warm-wet, or cool-wet) in year *t*+1, given climate state *x* (*e.g.*, cool-wet) in year *t* (Table S4).

From the simulations, we extracted the stochastic population growth rate (λ*_s_*), and population size projections (*n*) to evaluate the risks of future population decline and extinction. The long-term growth rate λ*_s_*, representing the average ratio of population sizes *n*(*t*+1)/*n*(*t*) (Ellner & Rees, 2007), was calculated across 100 iterations per simulation using the *lambda* function in ‘ipmr’. One hundred iterations were sufficient for λ*_s_* estimates to converge to their expected asymptotic behaviour (not shown). Additionally, a burn-in of 10 time steps was used to reduce the influence of transient dynamics on the analysis (Stott *et al*., 2011). For extinction risk analysis, plant densities (*n*) were extracted at each time step using the *collapse_pop_state* function in ‘ipmr’, and assigned a persistence score. Populations were assigned persistence scores as a binomial variable: 1 for time-steps where *n* > 0, and 0 otherwise.

### Hypothesis testing

To assess the effects of climate change and harvesting on population growth rate (H1-H5), we developed a multiple linear regression model using ln(λ*_s_*) as the response variable. The model included intercept terms for each climate scenario (*C_x_*) and harvesting scenario (θ*_p_*), with harvest frequency (θ*_f_*) as a continuous covariate to evaluate the thresholds of population decline. Based on AIC scores, a saturated model including all possible interaction terms provided the best fit. We estimated separate effects for harvest frequency dependent on climate scenario (*C_x_* × θ*_f_*) and harvesting scenario (θ*_p_* × θ*_f_*), as well as interactions between climate and harvesting scenario (θ*_p_* × *C_x_*), and three-way interactions (*C_x_* × θ*_p_* × θ*_f_*):

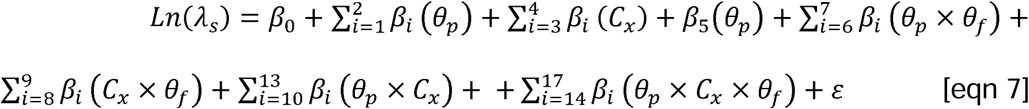

We used logistic regression to project extinction risk under climate change and harvesting. Since our IPM could not simulate the effects of carrying capacity on growing populations, we restricted this analysis to the 0-40 year time window to avoid unrealistic population projections. Additionally, complete separation in some predictors (*i.e.*, fitted probabilities of 1) was addressed by randomly switching 1% of persistence scores to 0. This step was implemented to mitigate standard error inflation with minimal influence on the model predictions. The adjusted model incorporated persistence scores as the response variable with a logit link function. Based on AIC comparisons, intercepts for *C_x_* and *θ_p_* were included, while year (*t*) and θ*_f_* were treated as continuous covariates. First and second order interaction terms were incorporated as follows:

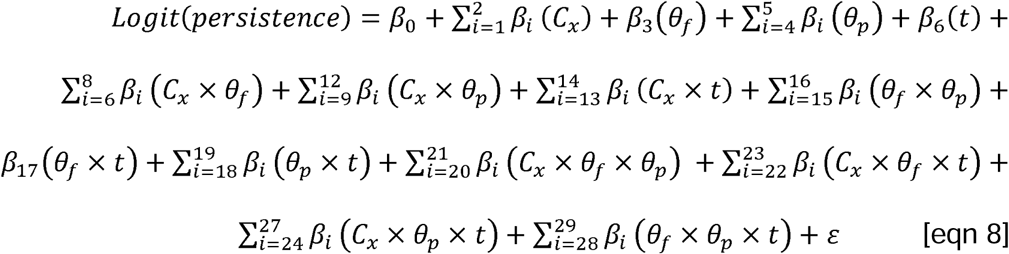

Our hypothesis testing was performed using Wald tests of estimated marginal means (EMMs) or trends (EMTs) using the ‘emmeans’ package (Lenth, 2023). Where appropriate, α levels were adjusted using Sidak corrections for multiple testing (Lenth, 2023). The tests were carried out as follows: H1: To test whether climate change impacts ln(λ*_s_*) independently of harvesting, we compared the EMMs of each climate change scenario against the EMM of the historical climate scenario using a left-tailed test. H2: To evaluate overall negative effects of harvest frequency, we tested whether the estimated marginal trend (EMT) of θ*_f_* was significantly less than zero (left-tailed test). H3: To assess interactions between climate change and θ*_f_,* we compared the EMTs of θ*_f_* for each climate scenario to the historical scenario using two-tailed tests. H4: To test whether plant harvesting has more severe effects than fruit harvesting, we contrasted the estimated marginal means of each plant harvesting scenario against the fruit harvesting scenario (left-tailed tests). H5: To test whether size-selective plant harvesting has more severe effects than non-selective harvesting, we performed EMM contrasts for the size-selective and non-selective harvesting scenarios (left-tailed tests).

### Perturbation analyses

To identify the demographic components that are vital for the persistence of *A. pearsonii* populations, we analysed the effects of perturbing each set of vital rate mean functions (σ, γ, τ, *φ_0_*, *φ_1_*, *φ_2_*, *φ_3_*, and *φ_4_*; Figure 2) on the stochastic population growth rate (λ*_s_*). Our analysis followed the function-level perturbation approach described in Griffith (2017), adapted for our stochastic IPM based on Ellner and Rees (2009) and Ellner *et al*. (2016). The effects of vital rate perturbations on λ*_s_* were quantified using elasticities, which represent the proportional change in λ*_s_* resulting from a proportional change in each vital rate function *f_t_*(*z*) or *f_t_*(*z*’,*z*) at size *z*. We chose elasticities over sensitivities because they allow for more direct comparisons across vital rates that are constrained between 0 and 1 (*e.g.*, survival) and those that are not (*e.g.*, reproduction; Zuidema and Franco, 2001).

For the perturbation analysis, we ran 1,000 iterations of our IPM, assuming a 0 harvest frequency and a historical climate scenario. From these iterations, we calculated λ*_s_* and extracted the time-series data for the left eigenvector (*v_t_*(*z*), reproductive value), right eigenvector (*w_t_*(*z*), population density), and the set of vital rate functions used in each iteration kernel *M*_t_(*z*,*z’*). The first 250 iterations of *v_t_*(*z*), *w_t_*(*z*), and *M_t_*(*z*,*z’*) were discarded from further calculations to eliminate transient dynamics (Stott *et al*., 2011).

The vital rate functions composing *M*_t_(*z*,*z’*) were used to formulate the perturbation kernel *C_t_*(*z,z’*) or function *C_t_*(*z*) sequences, which were identified by an appropriate substitution into the formulae defining each iteration kernel *M_t_*(*z,z’*) (Ellner and Rees, 2009). Perturbations of the vital rate kernels γ*(z,z’)* and *φ_4_(z,z’)* employed the proportional compensation method from Griffith (2017) to ensure that the integral of these kernels remained unchanged, while only altering pointwise probabilities.

Elasticities were calculated as:

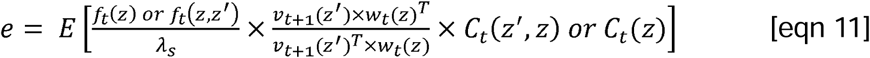

where the superscript *T* denotes the transpose of the corresponding left or right eigenvector. Elasticities were plotted as unidimensional functions (σ, τ, *φ_0_*, *φ_1_*, *φ_2_*, *φ_3_*,) or two-dimensional kernels (γ, *φ_4_*) to assess the effects of each vital rate at a given size/size transition on the stochastic population growth rate.

## Results

### Effects of climate change independent of harvesting (H1)

In the absence of harvesting, *A. pearsonii* populations face increased risk of decline under climate change relative to a historical climate scenario (Figure 3). Under hindcasted climates, our model predicted that at least 95% of all populations would remain viable when harvesting was absent (Figure 3). Under mild projected climate change (SSP126), the EMM of ln(λ*_s_*) was slightly, but significantly, lower than for the historical simulations (*P* < 0.001, Table 1). By contrast, the moderate climate change scenario (SSP245) projected a considerably stronger negative effect on the EMM of population growth (*P* < 0.001, Table 1), with ln(λ*_s_*) estimates falling below the threshold of decline for >50% of simulations when harvesting was absent (Figure 3).

**Figure 3.**
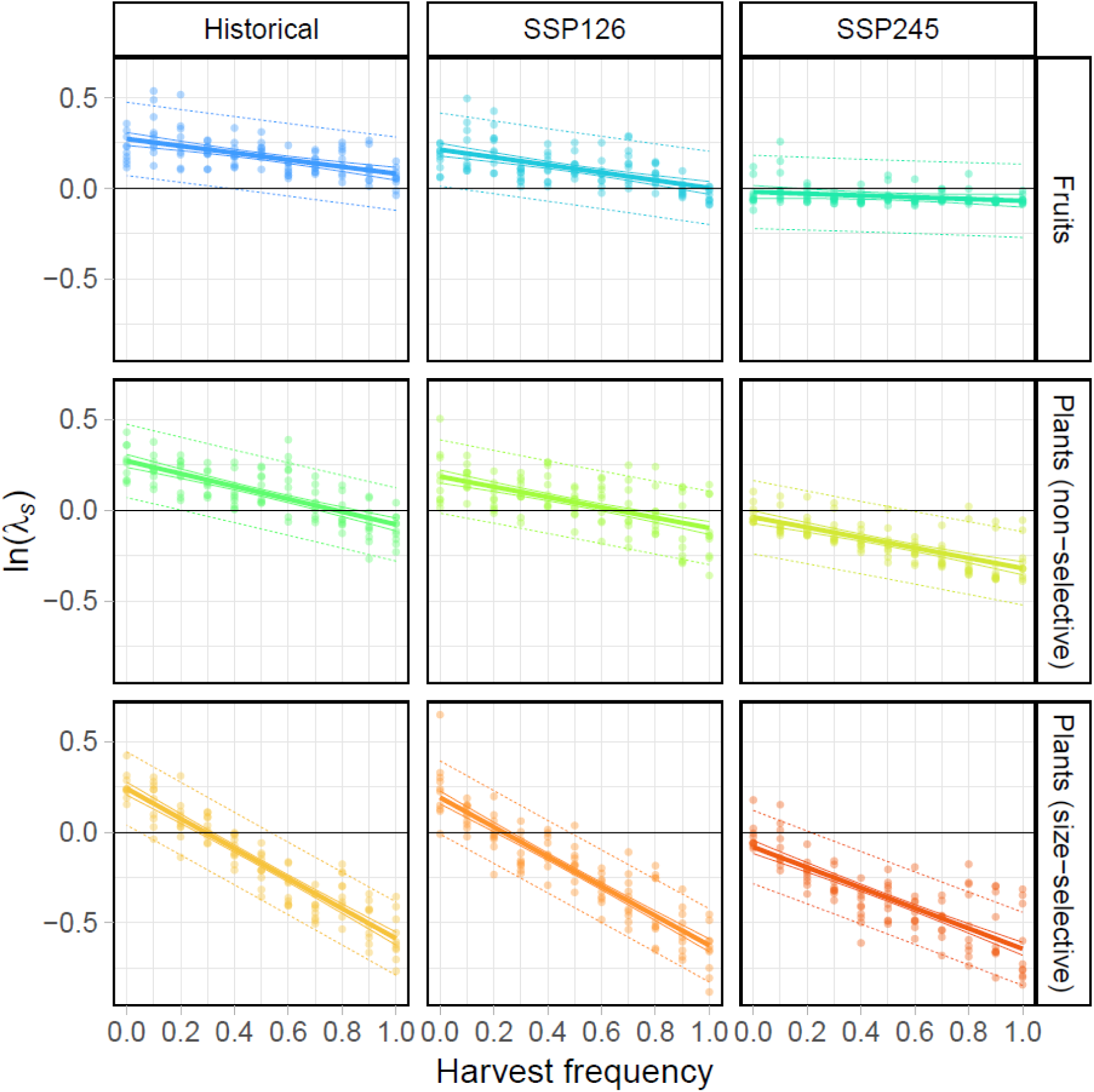
Climate change and harvesting lead to long-term population declines. Projected effects of climate change and harvesting on the long-term population growth rate (λ) of *Argyroderma pearsonii* as a function of harvest frequency. Fruit harvesting events encompassed the removal of 90% of fruit capsules from the population in a year (376 fruit capsules). Size-selective plant harvesting events encompassed the removal of 90% of individuals with a leaf pair larger than 10 mm in diameter (1,306 individuals). Non-selective plant harvesting events encompassed the removal of the same number of individuals from the population as in size-selective harvesting, except removal probability was independent of size. Points are ln(λ*_s_*) estimates for 100 iterations of our Integral Projection Model (*n* = 10 per climate scenario, harvesting scenario and harvesting frequency combination). Regression lines show fitted means (thick solid lines), with their 95% confidence intervals (thin solid lines), and 95% prediction intervals (dashed lines). The emboldened horizontal lines represent the thresholds for population growth (ln(λ*_s_*) > 0) and decline (ln(λ*_s_*) < 0).

**Table 1.**
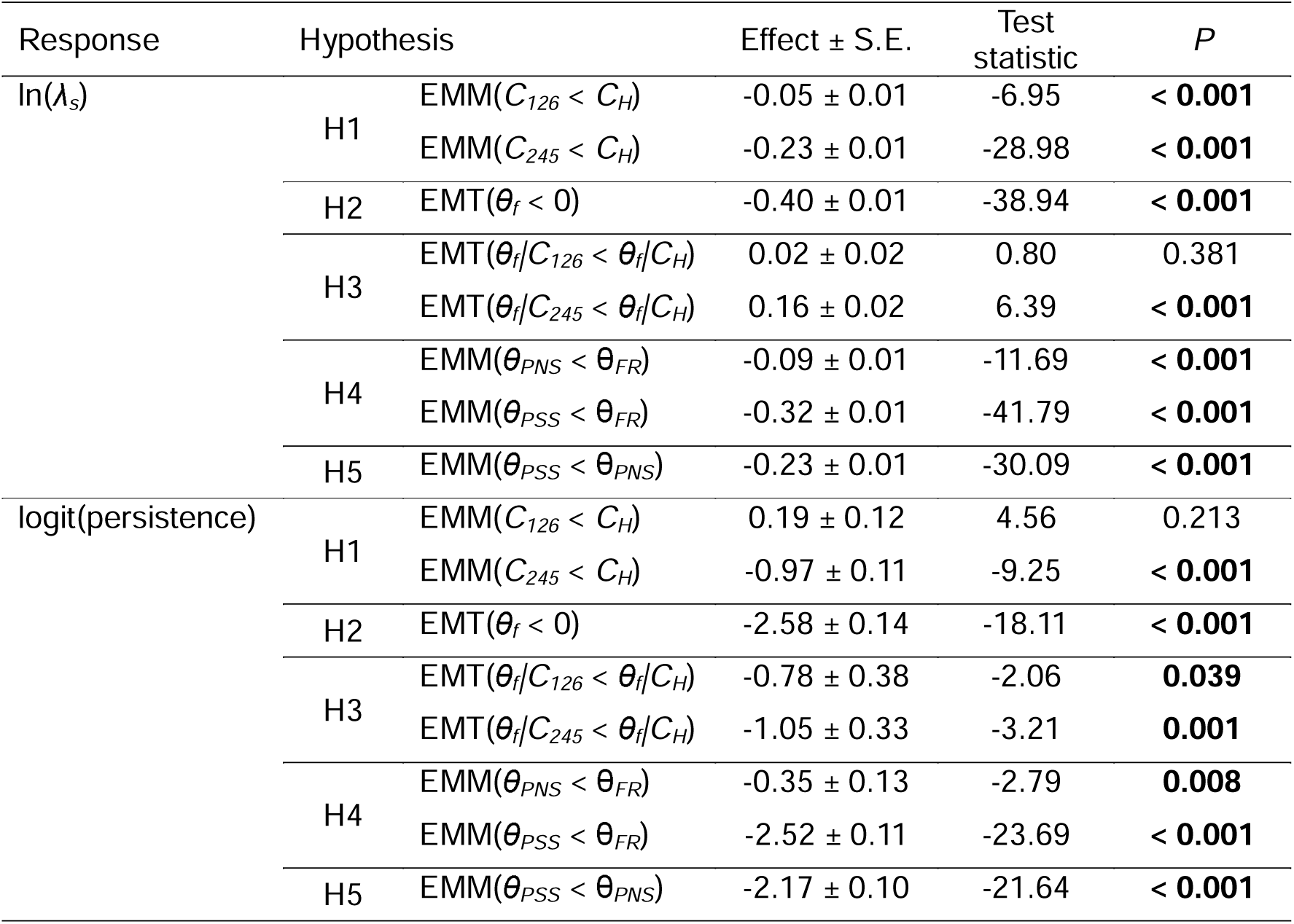
Results of hypothesis tests assessing the effects of climate scenario (*C_x_*), harvest scenario (*θ_p_*), and harvest frequency (*θ_f_*) on the growth rate (λ*_s_*) and persistence of *Argyroderma pearsonii* populations. Hypothesis tests were performed on the estimated marginal means (EMMs) or trends (EMTs) of log-linear (λ*_s_*) and logistic (persistence) models of IPM simulation outputs. Hypotheses are labelled according to their abbreviations in the text. Statistically significant results (α = 0.05) are indicated in bold. Subscript abbreviations for *C_x_* and *θ_p_* are as follows: H – hindcasted climates; 126 – mild (SSP126) climate change; 245 – moderate (SSP245) climate change; FR – fruit harvesting; PNS – non-selective plant harvesting; PSS – size-selective plant harvesting.

Despite increasing risks of long-term population decline, climate change did not increase the risk of short-term extinction in the absence of harvesting. Although our EMM analysis suggested that population persistence was significantly lower for SSP245 than for the historical scenario (*P* < 0.001, Table 1), zero out of 30 populations went extinct under SSP245 when harvest frequency was set to zero (Figure 4). Similarly, none of the replicate populations went extinct under SSP126, with the difference of this scenario’s EMM to that of the historical scenario being non-significant (*P* = 0.213, Table 1).

**Figure 4.**
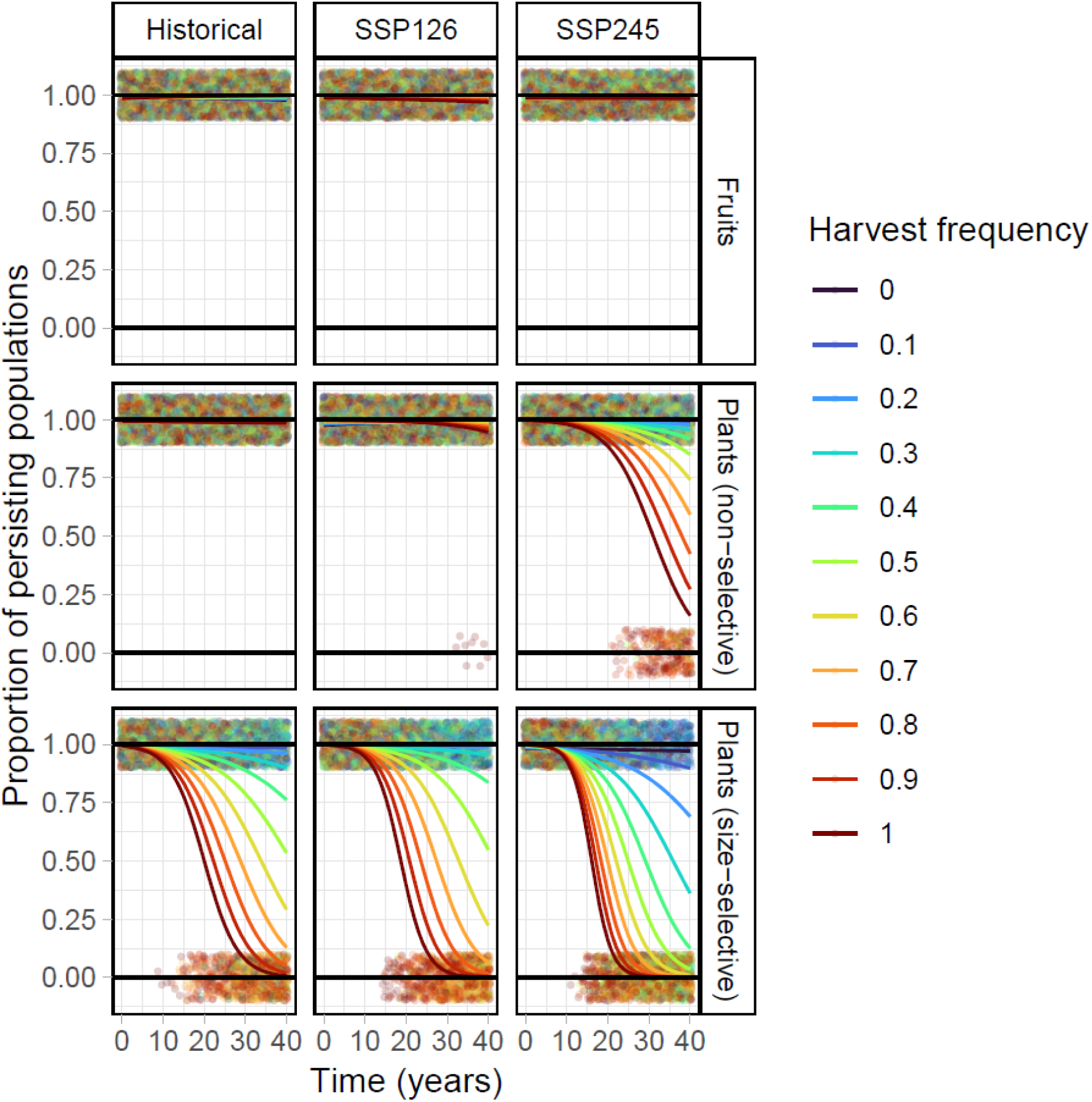
Plant harvesting increases short-term extinction risk. Projected extinction probability as a function of time for *Argyroderma pearsonii* under three climate scenarios and three harvesting scenarios with varying harvesting frequencies. Extinction scores were extracted from simulated population sizes (*i.e.*, surviving individuals · m^-2^) sourced from our Integral Projection Model (30 replicate simulations per combination of climate scenario, harvest scenario, and harvest frequency, with 40 time-steps per replicate). Populations were scored as extinct (points given a score of 1 in each panel) when the population size reached zero at a given time-step, and as surviving otherwise (points given a score of 0). Extinction probability was modelled as the fraction of extinct populations per replicate set using logistic regression.

### Effects of increasing harvest frequencies (H2)

Across all harvest types and climate scenarios, increases in the frequency of harvest events contributed to significant declines in the population growth rate (*P* < 0.001; Table 1; Figure 3). Assuming a historical climate scenario, our model predicted that increasing harvest frequencies from 0 to once every decade (*θ_f_* = 0.1) resulted in mean reductions in λ*_s_* from 1.30 to 1.12 for fruit harvesting, to 0.97 for non-selective plant harvesting, and to 0.62 for size-selective plant harvesting. The mean effect of harvest frequency was also significantly negative when considering short-term population persistence (Table 1, *P* < 0.001), with populations being more likely to go extinct in shorter timespans when plant harvesting frequencies increased (Figure 4).

### Interactions between harvest frequencies and climate change (H3)

Climate change was an important modulator of the effect of harvest frequency on population growth rate (*P* < 0.001, Table 1). However, a closer look at these effects reveals that, surprisingly, the slope of the SSP245 harvest frequency term was significantly greater than that of the historical climate scenario (*P* < 0.001), while the SSP126 harvest frequency term was not significantly different from that of the historical climate scenario (*P* = 0.380). Significance tests of the model coefficients indicated that the unexpected increase in the marginal trend of *C_SSP245_* was driven by the size-selective plant harvesting scenario (*β*(*C_SSP245_ × θ_plants (size-selective)_ × θ_f_*) effect ± S.E. = 0.12 ± 0.06, *t* = 2.02, *P* = 0.040) and not non-selective plant harvesting (effect ± S.E. = −0.07 ± 0.06, *t* = −1.21, *P* = 0.220). A possible reason for this result is that more frequent size-selective harvests drove λ*_s_* (a ratio of population size through time) towards its lower bound at zero, thus forcing the harvest frequency slope term to flatten.

The interaction between climate change and harvest frequencies had significant and more intuitive impacts on short-term extinction risk (*P* < 0.001, Table 1, Figure 4). Notably, the harvest frequency slope term of SSP245 was significantly less than for the historical scenario (*P* = 0.007), while the result for the SSP126 scenario was similar (*P* = 0.039). These results suggest that the short-term extinction risk associated with harvesting was amplified under climate change.

### Relative impacts of fruit harvesting and plant harvesting (H4 and H5)

Plant harvesting had more severe impacts on *A. pearsonii* populations than fruit harvesting (Figure 3, Figure 4). Size-selective, rather than non-selective, plant harvesting had the greatest impacts on population growth rates relative to fruit harvesting, with similar effects being evident when considering extinction risk (Table 1). Accordingly, size-selective plant harvesting had significantly stronger impacts than non-selective plant harvesting (*P* < 0.001).

### Perturbation analyses

Our perturbation analyses revealed that higher elasticity values were mostly concentrated above the 10 mm leaf size threshold that defines size-selective plant harvesting (Figure 5). Elasticity tended to be greater at larger sizes for all vital rates (Figure 5A, C, D), except the growth function (Figure 5B), with the latter indicating a stronger effect on population growth when smaller individuals shrank between time-steps. These results suggest that the survival and fecundity of larger individuals, and the potential for growth in early life stages, are crucial for the persistence of *A. pearsonii* populations. The importance of these larger individuals thus explains the relatively strong effects of size-selective plant harvesting (*i.e.*, removal of large plants and their reproductive contributions) compared to fruit and non-selective plant harvesting in the population simulations. Additionally, the population growth rate was elastic to perturbations of recruit sizes approaching the sizes typical of mature individuals (Figure 5D), suggesting that population regeneration with individuals of larger sizes may be expected to bolster population growth compared to regeneration with smaller individuals.

**Figure 5.**
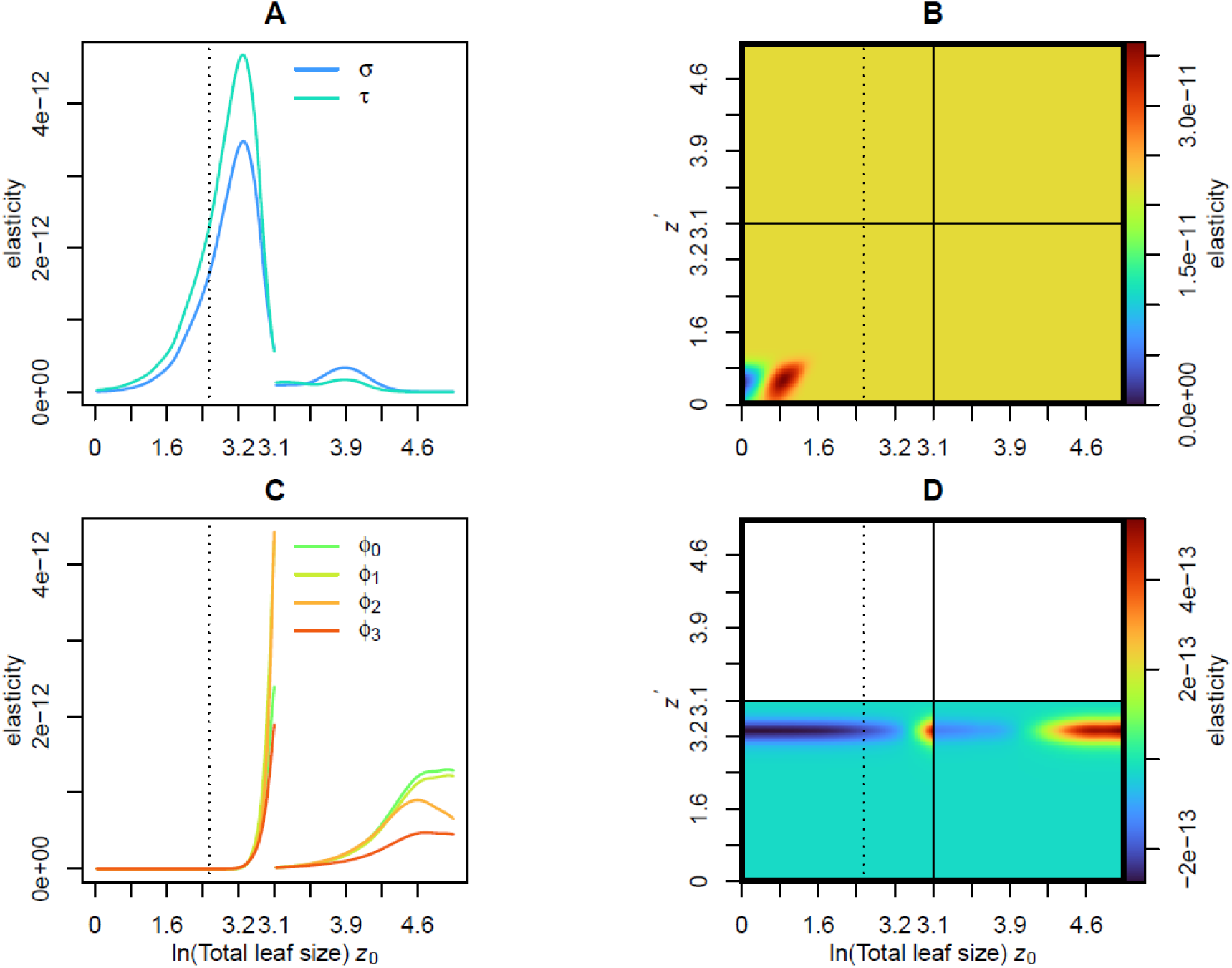
The population growth rate is most elastic to vital rate perturbations of the larger size classes. Stochastic kernel perturbation analyses for *Argyroderma pearsonii*, used to identify demographic components with the strongest influence on population growth rates. Perturbation analyses were based on a population simulation (1000 iterations) that assumed a historical climate scenario with a harvest frequency of zero. (A) Elasticity of the population growth rate λ to the survival (σ) and discrete state transition (τ, single-leaf or branched) functions. (B) Elasticity surface of the growth kernel (γ). (C) Sensitivity functions of the reproductive vital rates: flowering (*φ_0_*), fruit set (*φ_1_*), seed production (*φ_2_*), and recruit establishment (*φ_3_*). (D) Elasticity surface of the recruit size distribution (*φ*_4_). Vertical dotted lines in A-D represent the threshold minimum leaf size of 10 mm used for the size-selective plant harvesting simulations.

## Discussion

Global biodiversity faces growing threats from multiple anthropogenic pressures (Bellard *et al*., 2014), while forecasting the impacts of interacting threats is crucial for guiding conservation efforts (Co□té *et al*., 2016). Here, we built a stage-structured demographic model of the South African endemic succulent *Argyroderma pearsonii* and simulated the effects of climate change and illegal harvesting on population viability. In agreement with historical observations (Schmiedel *et al*., 2012), our model hindcasts viability for these populations under historical climate regimes, but forecasts important population declines, particularly when climate change interacts with illegal harvesting pressures. In the absence of harvesting, under mild (SSP126) or more severe climate scenarios (SSP245), populations are not expected to go extinct within 40 years. However, harvesting, especially when targeting large reproductive individuals, may significantly increase short-term extinction risk in conjunction with climate change. We also show that selective plant harvesting, rather than non-selective plant or fruit capsule harvesting, disproportionately affects population stability because it removes individuals that maintain population growth.

### Vulnerability to Climate Change

Increased drought and rising temperatures feature in several climatic projections for the Succulent Karoo (Hulme *et al*., 2001; MacKellar *et al*., 2007; Engelbrecht *et al*., 2015), while our population forecasts under different climate scenarios align with reported trends for dwarf Aizoaceae in the region (Musil *et al*., 2005, 2009; Midgley and Thuiller, 2007; Young *et al*., 2016; Milton *et al*., 2024; Grey and Atkinson, 2024). Musil *et al*. (2005) attributed mortality in quartz field succulent communities to reduced soil water potential resulting from experimentally increased ambient temperatures. Milton *et al*. (2024) attributed substantial (74%) population declines in the succulent *Bijlia dilatata* to failed recruitment and high mortality following two decades of drought (2002-2023) and rising temperatures in the Succulent Karoo. Grey and Atkinson (2024) reported increased mortality under drought conditions in *Argyroderma delaetti*, with the survival of juvenile individuals being disproportionately affected by drying compared to adults. Similarly, Midgley and Thuiller (2007) and Young *et al*. (2016) projected range contractions and possible extinction associated with future warming and drying trends in several dwarf Aizoaceae species.

Despite similarities in the outputs of our model to other ecological forecasts, future climate changes may pose greater risks to *A. pearsonii* than our model suggests. While our dataset describes demographic responses to extreme climatic conditions for the Knersvlakte under previously observed extremes (Schmiedel *et al*., 2012), these are far from the extremes of temperature and precipitation projected for the region by the UKESM global climate model (Mulcahy *et al*., 2022; Figure S4.1). Consequently, the impacts of future extremes have most likely not been captured in the underlying vital rates of our demographic model. Through open-top chamber experiments, Musil *et al*. (2005) showed that a 5.5□C increase in ambient temperatures resulted in a mortality rate increase of 53% for *A. pearsonii* relative to natural conditions. By contrast, our model projected a 10% decrease in average plant mortality under moderate climate change (*i.e.*, a 0.6□C increase, according to our climate classification dataset) relative to the historical climates. Currently, *A. pearsonii*’s conservation status is listed as least concern, according to the red list of South African plants (Burgoyne, 2006). With our relatively conservative findings in mind, we propose that this status be revised.

Recruitment success appears to be key for the viability of *A. pearsonii* populations under climate change. The UKESM model (Mulcahy *et al*., 2022) used to parameterise our simulations projected an increased frequency of warm, dry years in the future (Figure S4.2), while our IPM outputs suggest that this is correlated with declines in the population growth rate. Mechanistically, these findings make sense, as *A. pearsonii*, and many other dwarf Aizoaceae, produce many seeds (100s-1000s per fruit capsule - Duif, 2005; Jurgens and Witt, 2014), and depend on intense rainfall for seed dispersal and germination (Esler and Cowling, 1995; Parolin, 2006; Schmiedel *et al*., 2021; Eibes *et al*., 2022). Furthermore, warming disproportionately affects the survival of young individuals in the Aizoaceae (Musil *et al*., 2005; Milton *et al*., 2024; Grey and Atkinson, 2024). Thus, although only a small fraction of the total recruitment potential is usually realised (0.9% − 2.7% in our dataset, depending on year), these findings suggest that recruit numbers can be greatly enhanced through adequate precipitation inputs and cooler temperatures.

### Vulnerability to harvesting

Aside from records of illegally harvested plant seizures that indicate which species are targeted by harvesters, and crude estimates of reduction in population size (SANBI, 2024), there is little published information on the timing, frequency, and demographic targets of harvesting activities in the Succulent Karoo. This gap makes it difficult to simulate with certainty harvesting scenarios that are necessarily realistic and calls for more studies and harvesting data for reliable forecasts. Even though *A. pearsonii* is currently not the target of illegal harvest, our findings do provide insight into the likely population responses of targeted dwarf Aizoaceae (*e.g. Conophytum* and *Lithops*, SANBI, 2024). While specific responses to environmental change vary by species (Jurgens *et al*., 1999; Salguero-Gómez, 2017), several life history traits are shared among dwarf Aizoaceae suggesting broad applicability of our findings. Indeed, many dwarf Aizoaceae exhibit similarity in size-dependent survival and growth rates (Musil *et al*., 2005; Musker *et al*., 2020), adult life expectancy (>30 years, Schmiedel and Jurgens, 1999; Hartmann, 2004), seed traits (Liede and Hammer, 1990; Jurgens and Witt, 2014; Sukhorukov *et al*., 2023), and dispersal and germination strategies (Parolin, 2006; Schmiedel *et al*., 2021). Thus, given the ecological similarities and often limited geographic range of these species (Young *et al*., 2016; Eibes *et al*., 2022), our conclusions regarding *A. pearsonii* may extend to other endemic dwarf succulent Aizoaceae in South Africa’s quartz fields (Coutts *et al*., 2016).

We find that selective harvesting of mature plants poses the greatest threat to population viability, as it removes individuals with high reproductive value. Similar results have been found in an assessment of harvesting impacts on the population viability of *Aloe peglerae* (Aloaceae), which is targeted for sale as an ornamental plant (Pfab and Scholes, 2003). Available evidence indicates that large plants are the primary targets of illegal harvesting, while smaller plants and fruits are not targeted as frequently (Margulies, 2020). Similarly, larger individuals are preferred stock over smaller individuals among plant dealers, owing to their higher aesthetic value and tolerance of stress during transport and replanting (Margulies, 2020). Cultivated *Lithops* are not accepted for sale in European markets until they have grown to a size of at least 25 mm (Duif, 2005).

While there is considerable demand for seeds among nurseries (Duif, 2005), seed demand is smaller than for established plants because cultivation takes time and cannot immediately satisfy the demands of plant collectors (Margulies, 2020). Despite these challenges, our analyses suggest that, rather than plant harvesting, exclusive removal of fruit capsules would be the most sustainable harvesting scenario for dwarf succulents. Only when 90% of fruit were removed approximately every two years did our model project declines for more than 5% of simulated populations. This finding aligns well with a meta-analysis of sustainable seed harvesting practices in plants (Bucharova *et al*., 2024), which predicts that similar fractions of seeds may be harvested safely every two years for long-lived plants, such as *Argyroderma* and *Conophytum* (Schmiedel and Jurgens, 1999). However, the resilience of dwarf Aizoaceae to fruit harvesting practices may not be universal, as some species produce considerably lower numbers of seed per plant than *A. pearsonii* (Liede and Hammer, 1990; Duif, 2005; Jurgens and Witt, 2014).

In combination with climate change, selective harvesting of plants may impart stabilising selection for faster life history, such that plants reach sexual maturity at smaller sizes (Allendorf *et al*., 2008). Our perturbation analysis indicates that, while population growth rates are elastic to the establishment of recruits of various sizes, they are particularly elastic to the establishment of large recruits. Thus, on the one hand, we might expect selection for faster growth rates in the early years of life, since climate change is likely to disproportionately affect the survival probabilities of smaller individuals (Musil *et al*., 2005; Milton *et al*., 2024, Grey and Atkinson, 2024). On the other hand, selective harvesting of larger individuals and consequent removal of their reproductive outputs is expected to favour reproduction at earlier life stages. Shifts to earlier reproduction is a common pattern that has been revealed in studies of size-selective exploitation of wildlife (Hamon *et al*., 2000; Servanty *et al*., 2011). Coupled with the energetic and nutritional costs associated with flowering (Hartmann, 2004), this shift may drive adaptation towards a “live fast, die young” strategy, such that populations are better buffered against the removal of large individuals.

### Caveats and Limitations

Several meteorological factors not considered here likely influence demographic processes in dwarf succulents. Seasonal rainfall timing and the intensity of individual events are crucial for dispersal and recruitment (Parolin, 2006; Schmiedel *et al*., 2021), while fog and nocturnal dewfall may support survival and growth (Matimati *et al*., 2012). Although our model simplifies climate characterisation using annual averages, future studies should assess the effects of specific variables on vital rates within structured population models, potentially yielding more detailed insights (Ellner *et al*., 2016). Our datas*et al*.so lacks historical contingencies that affect population growth estimates (Schmiedel *et al*., 2012). For instance, flowering likely depends on lagged effects of water and nutrient availability (Hartmann, 2004). Expanding the dataset to include responses across varied climatic sequences could address this limitation. Finally, some projections indicated unrealistic population booms, likely due to unmodeled competition for limited germination sites (Hartmann, 2004) or initial patterns of transient amplification (Stott *et al*., 2011). Incorporating density-dependent factors may stabilise growth estimates under historical conditions. Notably, harvesting large, dominant plants might open recruitment sites, potentially buffering populations against decline (Martorell *et al*., 2015; Mitchell *et al*., 2021).

### Future Directions

Effective management of illegal wildlife trade requires a deeper understanding of how multiple threats shape population performance. In the case of some natural populations, these threats include harvesting, as well as the motivations behind exploitative activities (Hubschle and Margulies, 2023), and climate change. Structured population models are a useful tool for addressing such questions, as they can provide insight into the impacts of multiple threats, including human-wildlife conflicts (Davis, 2022), and the effective management of invasive alien species (Bogdan *et al*., 2021). However, critical knowledge gaps pertaining to the illegal plant trade include data on the frequency, intensity, and selectivity of harvesting events. These gaps make it challenging to accurately simulate realistic harvesting scenarios and highlight the need for additional studies and harvesting data to produce reliable forecasts. Collaboration between researchers and anti-poaching authorities is essential to monitor illegal activities and identify targeted species, while enhanced monitoring could generate valuable demographic datasets for threatened species. Additionally, systematic studies on the mechanisms driving population viability in threatened arid plant species are needed to better assimilate their life history traits and model their responses to anthropogenic disturbances (Salguero-Gómez *et al*., 2012).

### Conclusion

Using a mechanistic population model, we have shown that multiple anthropogenic disturbances are likely to compromise the resilience of succulent plant populations in the future. While both climate change and wild harvesting are expected to contribute to population declines in isolation, the impacts are compounded when both pressures act in unison, leading to significant extinction risks when plants are harvested from the wild. However, since fruit harvesting is less damaging than plant removal, nurseries could germinate seeds from wild-harvested fruit to supply the market with legally sourced plants. This strategy could reduce demand for poached plants and provide surplus stock for reintroduction efforts, supporting the recovery of threatened populations.

## Supporting information

SOM

## Acknowledgements

AE was supported by an Oxford-Africa Initiative grant (AFiOx-41) to RS-G and AGE, and the research was supported by a Newton Mobility grant (NMG\R2\170141) to AGE and RS-G. RS-G was supported by a NERC Pushing the Frontiers grant (NE/X013766/1). Fieldwork was supported by a Fulbright Grant to AGE.

## Notes

### Competing Interest Statement

The authors have declared no competing interest.

