## Supplementary material for "Poaching exacerbates the effects of climate change on the long-term viability of an endemic South African succulent plant species": SOM

Table of contents:

S1. Population sizes over the census period	P. 1

S2. Estimating seed production	P. 2

S3. Implementation of harvesting scenarios	P. 4

S4. Climatic data and continuity adjustments	P. 5

S5. Vital rate model selection	P. 9

**S1. Population sizes over the census period**

**Table S1. Population sizes over the census period.** See Fig. 1B for details of the locations of the six examined populations

| Population | 1999 | 2000 | 2001 | 2002 | 2003 | All years |
| --- | --- | --- | --- | --- | --- | --- |
| 1 | 376 | 269 | 686 | 958 | 407 | 1,411 |
| 2 | 404 | 206 | 735 | 635 | 290 | 1,190 |
| 3 | 235 | 262 | 374 | 365 | 336 | 472 |
| 4 | 261 | 358 | 749 | 949 | 327 | 1,340 |
| 5 | 52 | 47 | 49 | 54 | 45 | 84 |
| 6 | 208 | 114 | 130 | 119 | 95 | 279 |

**S2. Estimating seed production**

Seed production was estimated using a linear allometric equation predicting the number of seeds per fruit capsule by the maximum diameter of the fruit capsule (Linear model: no. of seeds = 283.72 × capsule diameter – 1394.2, *R^2^* = 0.66, *n* = 15, *P* < 0.05).

It was not possible to estimate seed production directly from the seed-capsule relationship in 1999 because capsule sizes had not been recorded for this year. We therefore simulated capsule sizes for the year 1999 using the capsule size and leaf size data for the years 2000-2003. To do this we formulated a cubic regression model with capsule size as the response and ln(total leaf size) as a continuous predictor. In addition, this model contained separate intercepts for whether individuals produced single or multiple capsules in a season (only 1 out of 337 fruiting individuals produced more than two capsules, Fig S1A). To capture heteroscedasticity in capsule sizes dependent on ln(total leaf size), we modelled the square root transformed residuals of the mean models as a quadratic function of ln(total leaf size) (Fig S1B). Following these regressions, capsule sizes for the year 1999 were simulated under a normal distribution. The mean and variance of the simulated capsule sizes were allowed to vary as a function of ln(total leaf size), parameterised by the coefficients of the mean and the back-transformed residual regressions respectively (Easterling *et al.*, 2000; Fig S1C). The correlation between actual and simulated capsule sizes for the years 2000-2003 using this approach was significant and moderately strong (Pearson’s *R* = 0.41; *P* < 0.001, Fig S1D). Seed output was then estimated for the simulated capsule sizes of year 1999, as for the years 2000-2003.

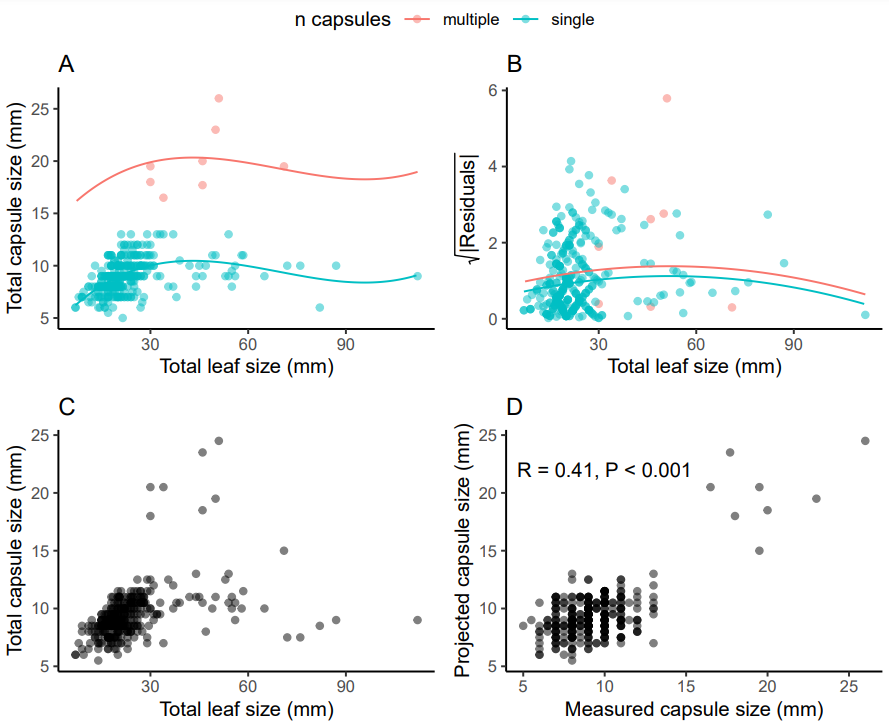

**Figure S2. Capsule sizes were projected for the year 1999 using the allometry of total capsule size and total leaf size for the years 2000-2003.** (A) Mean model of total capsule size predicted by total leaf size, fitted with a cubic slope term and separate intercepts for single/multiple capsules produced in a season. (B) Model of the residuals of (A) used to capture heteroscedasticity in the capsule size – leaf size relationship. (C) Projected total capsule sizes conditional on total leaf size, simulated under a normal distribution with the mean given by the predictions of model (A) and standard deviation given by the predictions of $\sqrt{\frac{\boldsymbol{\pi}}{\mathbf{2}}}$ × model (B)2. (D) Correlation of measured and projected capsules sizes.

**S3. Implementation of harvesting scenarios**

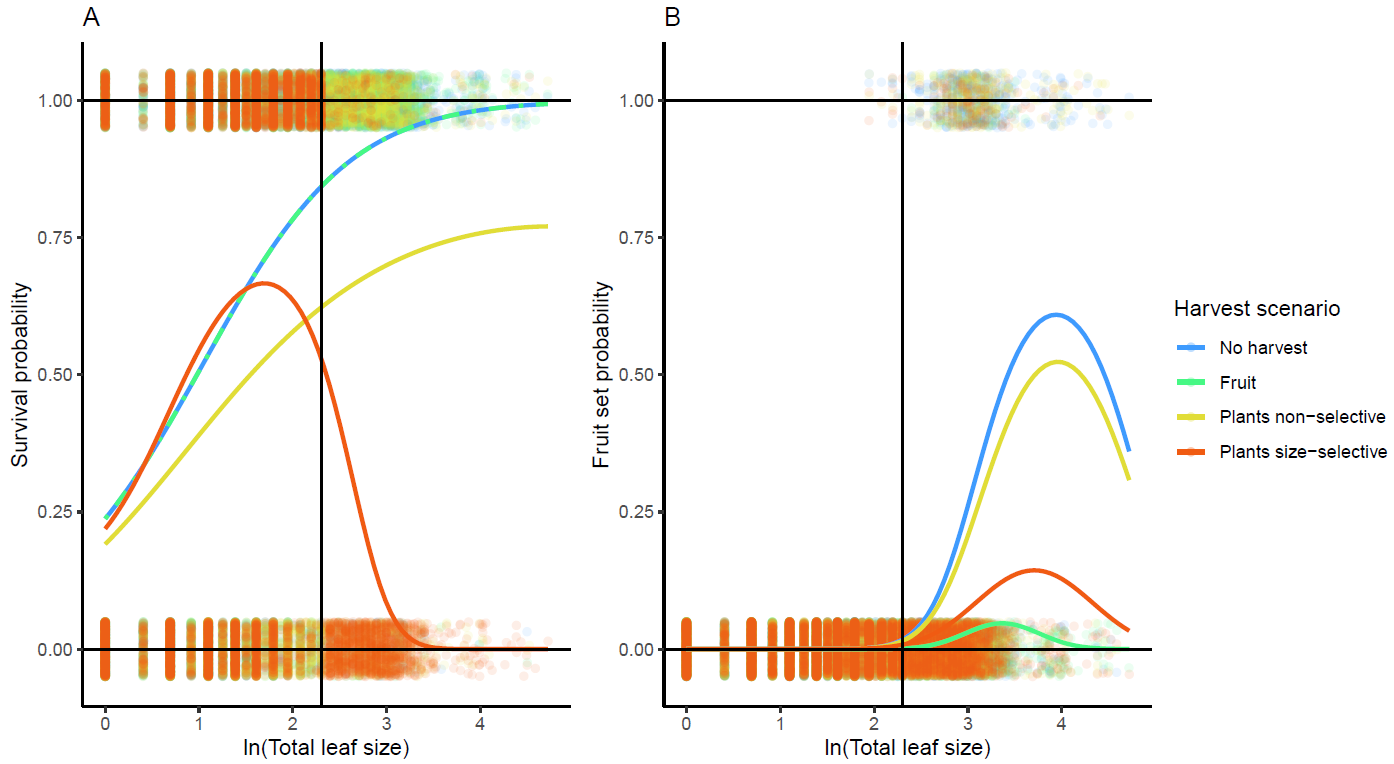

**Figure S3.** Effects of artificially defined harvesting scenarios on survival (A) and fruit set (B) as a function of the state variable ln(total leaf size). Coloured lines represent mean probabilities, estimated using cubic (A) and quadratic (B) polynomial regressions. Black vertical lines represent the threshold size for the size-selective plant harvesting scenario (10 mm).

**S4.** **Climatic data and continuity adjustments**

The historical and projected climate datasets exhibited two continuity issues, which were adjusted before proceeding with the construction of the Markovian climate matrices. First, the historical dataset comprised nine 0.9 km^2^ raster grid cells bounding the study region, while the projected datasets both comprised one 250 km^2^ grid cell. To account for these discrepancies in spatial resolution, we aggregated the historical dataset to the mean values of monthly temperature and precipitation across all nine grid cells. Thus, we sought to eliminate spatial variability in historic conditions, rather than risk spuriously increasing variance in projected conditions through statistical downscaling (Davy and Kusch, 2021; Kusch and Davy, 2022). Second, for the period of overlap between the two datasets (2016 – 2022), the projected climate dataset estimated monthly temperature and precipitation values that were respectively ~1⁰C warmer and ~8mm wetter than the historical dataset. To correct this bias, we adjusted the temperature and precipitation values of the projected datasets by subtracting the mean difference between the projected and historical datasets for these variables for the period of overlap. Following the continuity adjustments, we aggregated the monthly data to the mean annual temperature (MAT, *i.e.*, the mean of 12 monthly mean temperatures in ⁰C) and total annual precipitation (TAP, *i.e.,* the sum of precipitation over 12 months in mm) across the six sites. MAT and TAP represented the two axes of variation used to describe the climate of the study region each year.

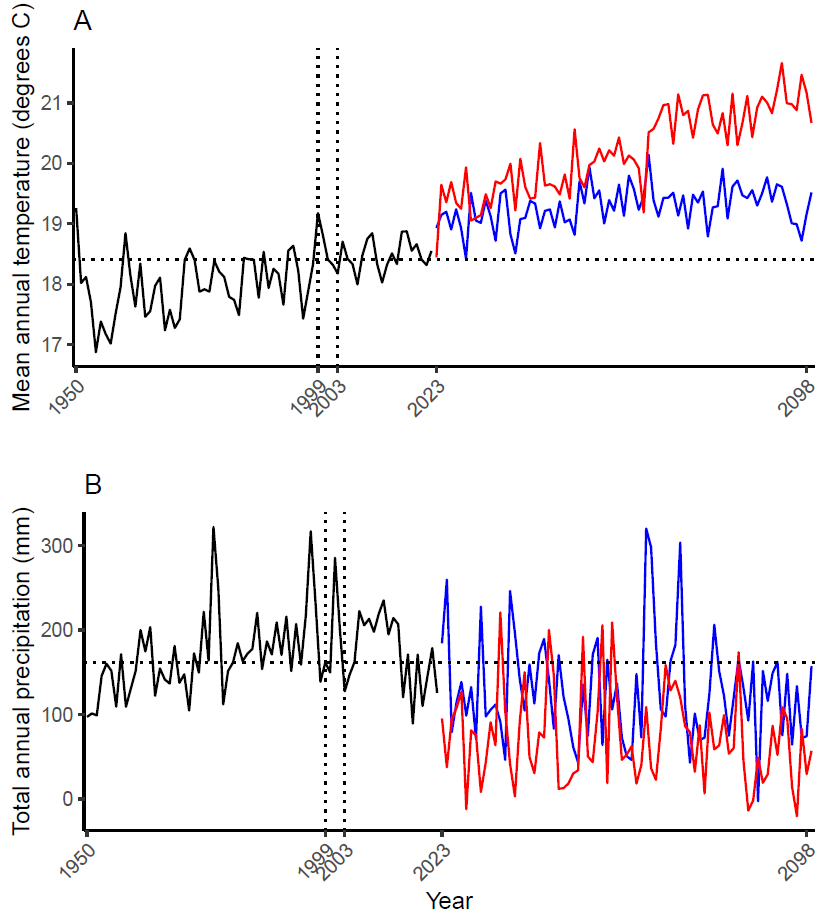

**Figure S4.1. Hindcasted (1950-2022) and projected (2023-2098) mean annual temperatures (A), and total annual precipitation (B) of the Knersvlakte bioregion of South Africa (31.4⁰S, 18.7⁰E).** Black lines are historical data acquired from the ERA 5-Land Reanalysis, and coloured lines are projected data from the UKESM1-0-LL Global Climate Model. Climate projections are according to two shared socioeconomic pathways (SSPs), representing mild (SSP126, blue), and moderate (SSP245, red) anthropogenic climate forcing. Vertical dotted lines indicate the census window used for the collection of demographic data for *Argyroderma pearsonii* (1999-2003). Horizontal dotted lines indicate the median values of each climatic variable for the census period, which were used to classify annual climates for the Markovian transition matrices.

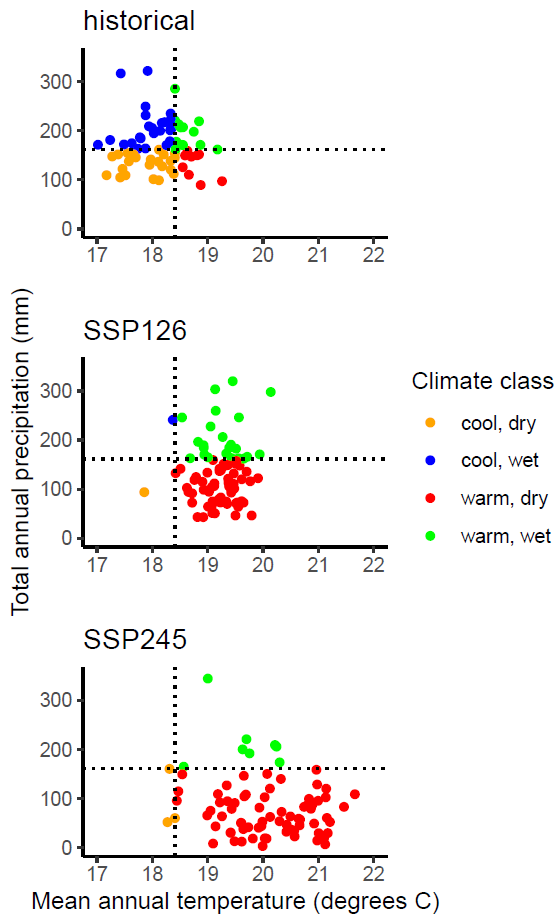

**Figure S4.2. Annual climate classifications (coloured points) used in the formulation of Markovian transition matrices under historical climates (years 1950-2022), and mild (SSP126), and moderate (SSP245), projected climate change (years 2023-2098).** Dotted lines are the median values of mean annual temperature and total annual precipitation of the five census years (1999-2003) used to define the thresholds for climate classification.

**Table S4. Markovian matrices used to simulate demographic transitions under three climate scenarios.** Cool-dry years were represented by vital rates estimated for the 2002-2003 transition; cool-wet years by 2001-2002; warm-dry by 1999-2000; and warm-wet by 2000-2001.

|  |  | Climate (*t*+1) | | | |
| --- | --- | --- | --- | --- | --- |
| Scenario | Climate (*t*) | Cool-dry | Cool-wet | Warm-dry | Warm-wet |
| Historical | Cool-dry | 0.54 | 0.25 | 0.08 | 0.13 |
|  | Cool-wet | 0.26 | 0.52 | 0.07 | 0.15 |
|  | Warm-dry | 0.22 | 0.44 | 0.00 | 0.33 |
|  | Warm-wet | 0.15 | 0.31 | 0.31 | 0.23 |
| SSP126 | Cool-dry | 0.00 | 0.00 | 1.00 | 0.00 |
|  | Cool-wet | 0.00 | 0.00 | 0.00 | 1.00 |
|  | Warm-dry | 0.02 | 0.00 | 0.75 | 0.24 |
|  | Warm-wet | 0.00 | 0.00 | 0.65 | 0.35 |
| SSP245 | Cool-dry | 0.00 | 0.00 | 0.67 | 0.33 |
|  | Cool-wet | 0.00 | 0.00 | 0.00 | 0.00 |
|  | Warm-dry | 0.03 | 0.00 | 0.89 | 0.08 |
|  | Warm-wet | 0.13 | 0.00 | 0.88 | 0.00 |

**S5.** **Vital rate model selection**

To optimise the explanatory power, goodness of fit, and parsimony of our overall IPM, candidate vital rate models for survival (*σ*), growth (*γ*), flowering (*φ*_0_), fruit set (*φ*_1_), and seed production (*φ*_2_) were subjected to a four-step model selection procedure. Each step of model selection addressed: (1) the value of considering multiple states for each vital rate, based on transitions between multiple leaf pairs and single leaf pairs in a season, (2) the best choice of state variable for branched (*i.e.* multiple leaf pair) individuals, and (3) model diagnostics.

1. *Single or multiple states*

In the initial round of model selection, we considered the alternative possibilities of defining either a single state or two-state IPM, based on the occurrence of branching. Here, we used total leaf size as the continuous state variable predictor for each vital rate. However, as total leaf sizes were strongly skewed toward smaller individuals (Figure S5), we also considered the use of ln-transformed values of total leaf size. For the single state IPM option, the vital rates of all individuals were treated equivalently, regardless of whether an individual had one or multiple leaf pairs. For the two-state option, the vital rates of individuals were treated differently based on whether they had one leaf pair or had branched to form multiple leaf pairs.

Branching implicated a need to consider multiple continuous states in our analyses, as our data suggest that multiple leaf pair individuals can produce multiple fruit capsules in a season (Figure 1A), while single leaf pair individuals cannot. Thus, branching was expected to have a considerable impact on individual reproduction, assuming that the seed number by capsule size relationship was similar among branched and single-pair individuals (Figure S2). To evaluate the effect of branching on vital rates, we compared vital rate models with (1) either of the two continuous state variable candidates as the sole predictor of each vital rate; (2) models with additional intercept terms for single/multiple leaf pair individuals; and (3) models with additional intercept and slope terms for single/multiple leaf pair individuals. Additionally, we considered linear, quadratic, and cubic polynomial terms of each state variable to capture complexities in the relationships between vital rates and state variables in the initial model selection procedure.

The best fitting vital rate models were identified from those nested within a saturated generalised linear mixed model structure, implemented in the package glmmTMB (Brookes *et al.*, 2017). The saturated model contained a random intercept term for site, the fixed effects of year (1999 – 2002), one of the candidate state variables (total leaf size or ln(total leaf size)) with a cubic polynomial term, leaf pair quantity (single or multiple leaf pairs), and a polynomial interaction between leaf pair quantity and the state variable. Nesting of candidate models proceeded with the removal of the state × leaf pair quantity interaction, the main effect of leaf pair quantity and reduction of the polynomial term to quadratic and linear effects. Effects that were retained in all the models included the random intercept site term and year. Owing to the simulation of capsule size values based on ln(total leaf size) in the year 1999 (Above), the appraisal of seed number vital rate models was restricted to the years 2000-2002, to avoid biasing model selection with the implicit dependency of capsule sizes in 1999 on ln(total leaf size). Additionally, the seed production model included a categorical predictor for whether an individual produced one or multiple fruit capsules in a year. As with the year and site intercepts, the single/multiple capsule predictor was also retained among all the candidate models for seed production. The comparison of nested models was based on Akaike’s Information Criterion (AIC) for all vital rates, while maximum likelihood pseudo-*R*^2^ was used for comparing growth models with different response variables (*e.g.*, ln(total leaf size) *vs*. total leaf size responses).

The results of the initial round of model selection indicated that (1) ln transformed total leaf sizes, and (2) leaf pair quantity and/or leaf pair quantity × ln(total leaf size) interactions had the best fit to the data (Table S5.1). Therefore, we opted for a two-stage model for single/multiple leaf pairs and eliminated raw total leaf size values as a candidate state variable. Since the only two state variables that were available in our dataset for single pair individuals were total leaf size and ln(total leaf size), we modelled the demographic transitions of single pair individuals using ln(total leaf size) in our IPM.

2. *State variable for branched individuals*

Our demographic dataset contained an additional state variable – plant size – for multiple leaf pair individuals, which was considered as a state variable for our IPM alongside total leaf size. Plant size was defined as the maximum diameter across the exposed portion of the plant, measured in mm.

For our second round of model selection, we compared the explanatory power of the ln(total leaf size), plant size and ln(total plant size) as state variable predictors of each vital rate in branched individuals using the same AIC and pseudo-*R*^2^ approach as before. The results of this round of model selection indicated that the use of ln(total leaf size) as a state variable produced similar goodness of fit results to plant size in most instances, while ln(Plant size) usually had worse fit than either of the other two candidate state variables (Table S5.2). However, the use of plant size as the state variable often resulted in model convergence failure. Therefore, for the sake of stability in further model fitting, and consistency with the choice of state variable in single-pair individuals, we decided to use ln(total leaf size) as the state variable for subsequent fitting of all vital rate models.

3. *Model diagnostics*

Following the state variable selection procedure, the selected models were subjected to residual simulation diagnostics using the ‘DHARMa’ package (Hartig, 2018). The model diagnostics included quantile-quantile plots of the simulated residuals, Kolmogorov-Smirnov tests to assess the appropriateness of the error distribution, dispersion tests, and outlier tests. Homogeneity of variance was assessed using Quantile Generalised Additive Models (QGAMs) fitted to the standardised residuals for continuous predictors, and Levene tests for categorical predictors. This approach was followed by posterior predictive checks and multicollinearity checks using the ‘performance’ package (Lüdecke *et al.*, 2021). If any issues with the fit were detected, the model structures were adjusted accordingly, and the diagnostic procedure was repeated on the newly formulated models until the fit could not be improved any further. All newly formulated models were compared to the originally selected models with AIC. Where heteroscedasticity was evident (*i.e.*, in growth models), the residuals were subjected to a regression against the state variable. Diagnostics for the residual regressions were performed as for the mean models. All updated models were compared to the original models with AIC.

While the initial rounds of model selection indicated that growth was best predicted by polynomial (*i.e.*, cubic and quadratic) fits, residual patterns suggested a more complex growth relationship that could not be captured adequately using polynomial regression. Therefore, the growth of both single and branched individuals was described using generalised additive models (GAMs, Wood, 2017), implemented in the ‘mgcv’ package (Wood, 2003). Thin plate penalised regression splines were used to fit the relationship between ln(total leaf sizes) in times *t*+1 and *t*, with a parametric term included for year and a random effect smooth term for site. The model for single-pair individuals used a spline with ten basis functions and assumed a scaled-*t* error distribution to account for overdispersion in the residuals, while the model for branched individuals used a spline with four basis functions and assumed a Gaussian error distribution. The site smooth term used one basis function for each site (*i.e.* six basis functions).

**Table S5.1. Results of Generalized Linear Mixed Model selection experiments evaluating the importance of state variable (Total leaf size or ln(Total leaf size)), number of leaf pairs (single or multiple) and interactions between the state variable and the number of leaf pairs for the goodness-of-fit of the vital rate regressions (σ = survival probability, μγ = growth mean, φ_0_ = flowering probability, φ_1_ = fruit set probability, φ_2_ = seed production).** Model comparisons were performed with AICc and maximum likelihood pseudo-R^2^ (the latter for comparison of growth mean *μγ* models with different response variables). The degree column refers to the polynomial degree of the continuous state variable (1 = linear, 2 = quadratic, 3 = cubic). Cells with darker green shading indicate better fitting models according to the AICc and pseudo-R^2^ statistics. Blank cells indicate that a model failed to converge or was otherwise deemed unsuitable for further analyses.

| State | Degree | Leaf pairs | State × Leaf pairs | *σ* | *μγ* | | *φ_0_* | *φ_1_* | *φ_2_* |
| --- | --- | --- | --- | --- | --- | --- | --- | --- | --- |
|  |  |  |  | AICc | AICc | pseudo-*R*^2^ | AICc | AICc | AICc |
| ln(Total leaf size) | 1 | X | X | 8366.51 | -2348.88 | 0.93 | 1926.86 | 331.69 | 49416.13 |
| ln(Total leaf size) | 1 | X |  | 8365.62 | -2349.68 | 0.93 | 1939.98 | 330.09 | 52175.60 |
| ln(Total leaf size) | 1 |  |  | 8365.46 | -2347.67 | 0.93 | 2000.64 | 329.46 | 56454.93 |
| ln(Total leaf size) | 2 | X | X | 8361.03 | -2704.02 | 0.93 | 1921.61 |  | 47971.97 |
| ln(Total leaf size) | 2 | X |  | 8357.84 | -2686.86 | 0.93 | 1917.62 | 331.96 | 48872.55 |
| ln(Total leaf size) | 2 |  |  | 8355.99 | -2561.58 | 0.93 | 1924.15 | 330.68 | 50472.16 |
| ln(Total leaf size) | 3 | X | X | 8364.80 | -2931.78 | 0.93 | 1924.51 | 334.73 | 47820.34 |
| ln(Total leaf size) | 3 | X |  | 8359.28 | -2857.60 | 0.93 | 1919.45 | 333.54 | 48315.58 |
| ln(Total leaf size) | 3 |  |  | 8357.29 | -2859.46 | 0.93 | 1926.14 | 331.81 | 50355.14 |
| Total leaf size | 1 | X | X | 8772.41 | 24337.61 | 0.91 | 2015.78 | 332.36 | 50209.91 |
| Total leaf size | 1 | X |  | 8800.61 | 24443.01 | 0.91 | 2155.25 | 330.42 | 57996.74 |
| Total leaf size | 1 |  |  | 8823.40 | 24778.95 | 0.91 | 2316.82 | 329.29 | 58967.61 |
| Total leaf size | 2 | X | X | 8532.95 | 24145.60 | 0.92 | 1925.29 | 332.26 | 48726.78 |
| Total leaf size | 2 | X |  | 8702.73 | 24271.60 | 0.92 | 1988.54 | 331.90 | 49445.69 |
| Total leaf size | 2 |  |  | 8700.81 | 24352.60 | 0.91 | 2056.27 | 331.16 | 53388.37 |
| Total leaf size | 3 | X | X | 8401.18 | 23928.89 | 0.92 |  | 334.14 |  |
| Total leaf size | 3 | X |  | 8511.53 | 24240.35 | 0.92 | 1941.53 |  |  |
| Total leaf size | 3 |  |  | 8547.45 | 24252.87 | 0.92 | 1946.83 |  |  |

**Table S5.1. Results of Generalized Linear Mixed Model selection experiments evaluating the importance of state variable (ln(Total leaf size), Plant size, or ln(Plant size)) for the goodness-of-fit of the vital rate regressions for multiple leaf pair individuals.** Further details as for Table S5.1.

| Leaf pairs | State | Degree | *σ* | *μγ* | | *φ_0_* | *φ_1_* | *φ_2_* |
| --- | --- | --- | --- | --- | --- | --- | --- | --- |
|  |  |  | AICc | AICc | Pseudo-*R*^2^ | AICc | AICc | AICc |
| single | ln(Total leaf size) | 1 | 8339.4 | -2543.45 | 0.924 | 1793.95 | 285.65 | 38848.1 |
| single | ln(Total leaf size) | 2 | 8332.1 | -2903.93 | 0.929 | 1788.3 | 285.2 | 38808.9 |
| single | ln(Total leaf size) | 3 | 8333.9 | -3138.81 | 0.933 | 1789.55 | 287.2 | 38547.5 |
| multiple | ln(Total leaf size) | 1 | 33.77 | 77.64 | 0.405 | 128.87 | 57.17 | 4341.72 |
| multiple | ln(Total leaf size) | 2 |  | 75.94 | 0.44 | 130.97 |  | 3726.13 |
| multiple | ln(Total leaf size) | 3 |  |  |  | 132.39 | 58.03 | 3717.42 |
| multiple | Plant size | 1 | 33.79 | 351.92 | 0.854 | 129.2 | 51.89 | 4401.54 |
| multiple | Plant size | 2 | 36.02 | 354.88 | 0.853 | 131.6 | 53.56 | 4397.41 |
| multiple | Plant size | 3 |  | 353.81 | 0.863 | 129.75 |  | 4300.72 |
| multiple | ln(Plant size) | 1 | 33.74 | -49.02 | 0.807 | 129.11 | 51.46 | 4405.39 |
| multiple | ln(Plant size) | 2 |  | -61.93 | 0.853 | 131.16 | 53.04 | 4402.75 |
| multiple | ln(Plant size) | 3 |  | -62.93 | 0.863 | 132.07 | 55.72 | 4395 |

**Table S5.3. Vital rate model structures used to parameterize an Integral Projection Model for *Argyroderma pearsonii* to examine the effects of harvesting and climate change.** The vital rate models are identified by site*_j_* (*j* ϵ [1;6]), capsule quantity *ω* (single or multiple, see $\varphi_{2_{b}}$), census year *t_y_* (*y* ϵ [1999;2002]), and harvesting scenario *θ_p_* (*p* ϵ [non-selective-plant, size-selective-plant, and fruit harvesting]). The state variable, *z*, is substituted by *s* and *b* for each possible state transition (*z*,*z*’), where *s* refers to individuals with a single leaf pair and *b* refers to individuals with multiple leaf pairs. Model type abbreviations are as follows: GAM = Generalised Additive Model; GLM = Generalised Linear Model; GLMM = Generalised Linear Mixed Model; LM = Linear Model.

| Vital rate | Model equation |
| --- | --- |
| Survival single stasis (binomial GLMM) | $Logit\left( \sigma_{s,s^{'}} \right)=\beta_{0}+\sum_{i=1}^{3} \beta_{i}\left( \theta_{p} \right)+\sum_{i=4}^{6} \beta_{i}\left( t_{y} \right)+\beta_{7}\left( s \right)+\beta_{8}\left( s^{2} \right)+\sum_{i=9}^{11} \beta_{i}\left( \theta_{p}\times s \right)+\sum_{i=12}^{14} \beta_{i}\left( \theta_{p}\times s^{2} \right)+\sum_{i=15}^{17} \beta_{i}\left( t_{y}\times s \right)+\sum_{i=18}^{20} \beta_{i}\left( t_{y}\times s^{2} \right)+\sum_{i=21}^{29} \beta_{i}\left( {\theta_{p}\times t}_{y}\times s \right)+\sum_{i=30}^{38} \beta_{i}\left( \theta_{p}\times t_{y}\times s^{2} \right)+\sum_{i=1}^{6} b_{i}\left( {site}_{j} \right)+\varepsilon$ |
| Survival branching (binomial GLMM) | $Logit(\sigma_{s,b'})=\beta_{0}+\sum_{i=1}^{3} \beta_{i}(\theta_{p})+\sum_{i=4}^{6} \beta_{i}(t_{y})+\beta_{7}(s)+\sum_{i=1}^{6} b_{i}({site}_{j})+\varepsilon$ |
| Survival branched stasis (binomial GLM) | $Logit(\sigma_{b,b'})=\beta_{0}+\sum_{i=1}^{3} \beta_{i}(\theta_{p})+\sum_{i=4}^{6} \beta_{i}(t_{y})+\varepsilon$ |
| Survival dieback (binomial GLM) | $Logit(\sigma_{b,b'})=\beta_{0}+\sum_{i=1}^{3} \beta_{i}(\theta_{p})+\sum_{i=4}^{6} \beta_{i}(t_{y})+\varepsilon$ |
| Mean growth single stasis (scaled-*t* GAM) | $\mu\gamma_{s,s'}=\beta_{0}+\sum_{i=1}^{3} \beta_{i}\left( t_{y} \right)+f\left( s,t_{y} \right)+\sum_{i=1}^{6} b_{i}({site}_{j})+\varepsilon$ |
| Variance in growth single stasis (scaled-*t* GAM) | $Ln(\left\vert\varepsilon\gamma_{s,s^{'}} \right\vert)=\beta_{0}+f\left( s \right)+\varepsilon$ |
| Growth distribution single stasis (Gaussian density function) | $\gamma_{s,s'}=\mathrm{Normal}(s',\mu\gamma_{s,s^{'}}(t),\sqrt{\frac{\pi}{2}}.e^{\left\vert\varepsilon\gamma_{s,s^{'}} \right\vert}$) |
| Mean growth branching (scaled-t GAM) | $\mu\gamma_{s,b'}=\beta_{0}+\sum_{i=1}^{3} \beta_{i}\left( t_{y} \right)+f\left( s,t_{y} \right)+\sum_{i=1}^{6} b_{i}({site}_{j})+\varepsilon$ |
| Variance in growth branching (scaled-t GAM) | $Ln(\left\vert\varepsilon\gamma_{s,b^{'}} \right\vert)=\beta_{0}+f\left( s \right)+\varepsilon$ |
| Growth distribution branching (Gaussian density function) | $\gamma_{s,b'}=\mathrm{Normal}(b',\mu\gamma_{s,b^{'}}(t),\sqrt{\frac{\pi}{2}}.e^{\left\vert\varepsilon\gamma_{s,b^{'}} \right\vert}$) |
| Mean growth branched (Gaussian GAM) | $\mu\gamma_{b,b'}=\beta_{0}+\sum_{i=1}^{3} \beta_{i}\left( t_{y} \right)+f\left( b \right)+\varepsilon$ |
| Variance in growth branched (Gaussian GAM) | $Ln(\left\vert\varepsilon\gamma_{b,b^{'}} \right\vert)=\beta_{0}+f\left( b \right)+\varepsilon$ |
| Growth distribution branched (Gaussian density function) | $\gamma_{b,b'}=\mathrm{Normal}(b',\mu\gamma_{b,b^{'}}(t),\sqrt{\frac{\pi}{2}}.e^{\left\vert\varepsilon\gamma_{b,b^{'}} \right\vert}$) |
| Mean growth dieback (Gaussian GAM) | $\mu\gamma_{b,s'}=\beta_{0}+\sum_{i=1}^{3} \beta_{i}\left( t_{y} \right)+f\left( b \right)+\varepsilon$ |
| Variance in growth dieback (Gaussian GAM) | $Ln(\left\vert\varepsilon\gamma_{b,s^{'}} \right\vert)=\beta_{0}+f\left( b \right)+\varepsilon$ |
| Growth distribution dieback (Gaussian density function) | $\gamma_{b,s'}=\mathrm{Normal}(s',\mu\gamma_{b,s^{'}}(t),\sqrt{\frac{\pi}{2}}.e^{\left\vert\varepsilon\gamma_{b,s^{'}} \right\vert}$) |
| Flowering probability single (binomial GLMM) | $Logit\left( {\varphi0}_{s} \right)=\beta_{0}+\sum_{i=1}^{3} \beta_{i}\left( t_{y} \right)+\beta_{4}\left( s \right)+\beta_{5}\left( s^{2} \right)+\sum_{i=6}^{8} \beta_{i}(t_{y}\times s)+\sum_{i=9}^{11} \beta_{i}(t_{y}\times s^{2})+\sum_{i=1}^{6} b_{i}({site}_{j})+\varepsilon$ |
| Flowering probability branched (binomial GLM) | $Logit\left( {\varphi0}_{b} \right)=\beta_{0}+\sum_{i=1}^{3} \beta_{i}\left( t_{y} \right)+\beta_{4}\left( b \right)+\varepsilon$ |
| Fruit set probability single (binomial GLMM) | $Logit({\varphi1}_{s})=\beta_{0}+\sum_{i=1}^{3} \beta_{i}(\theta_{p})+\sum_{i=4}^{6} \beta_{i}(t_{y})+\beta_{7}(s)+\sum_{i=8}^{10} \beta_{i}(\theta_{p}\times s)+\sum_{i=11}^{13} \beta_{i}(t_{y}\times s)+\sum_{i=14}^{22} \beta_{i}(\theta_{p}\times t_{y}\times s)+\sum_{i=1}^{6} b_{i}({site}_{j})+\varepsilon$ |
| Fruit set probability branched (binomial GLM) | $Logit({\varphi1}_{b})=\beta_{0}+\sum_{i=1}^{3} \beta_{i}(\theta_{p})+\sum_{i=4}^{6} \beta_{i}(t_{y})+\beta_{7}(b)+\sum_{i=8}^{10} \beta_{i}(\theta_{p}\times b)+\varepsilon$ |
| Seed production single (Negative binomial GAM) | $Log\left( {\varphi2}_{s} \right)= \beta_{0}+\sum_{i=1}^{3} \beta_{i}\left( t_{y} \right)+f\left( s \right)+\sum_{i=1}^{6} b_{i}({site}_{j})+\varepsilon$ |
| Seed production branched (Negative binomial GLM) | $Log\left( {\varphi2}_{b} \right)= \beta_{0}+\sum_{i=1}^{3} \beta_{i}\left( t_{y} \right)+{\beta_{4}\left( \omega\right)+\beta}_{5}\left( s \right)+\beta_{6}\left( s^{2} \right)+\sum_{i=1}^{6} b_{i}({site}_{j})+\varepsilon$ |
| Recruitment probability (Beta GLMM) | $Logit\left( \varphi3 \right)= \beta_{0}+\sum_{i=1}^{3} \beta_{i}\left( t_{y} \right)+\sum_{i=1}^{6} b_{i}({site}_{j})+\varepsilon$ |
| Mean recruit size (Truncated Gaussian GLMM) | $\mu\varphi4= \beta_{0}+\sum_{i=1}^{3} \beta_{i}\left( t_{y} \right)+\sum_{i=1}^{6} b_{i}({site}_{j})+\varepsilon$ |
| Variance in recruit size (LM) | $Ln\left( \left\vert\varepsilon\varphi4 \right\vert\right)= \beta_{0}+\varepsilon$ |
| Recruit size distribution (Gaussian density function) | $\varphi4=\mathrm{Normal}(s',\mu\varphi4(t),\sqrt{\frac{\pi}{2}}.e^{\left\vert\varepsilon\varphi4 \right\vert}$) |

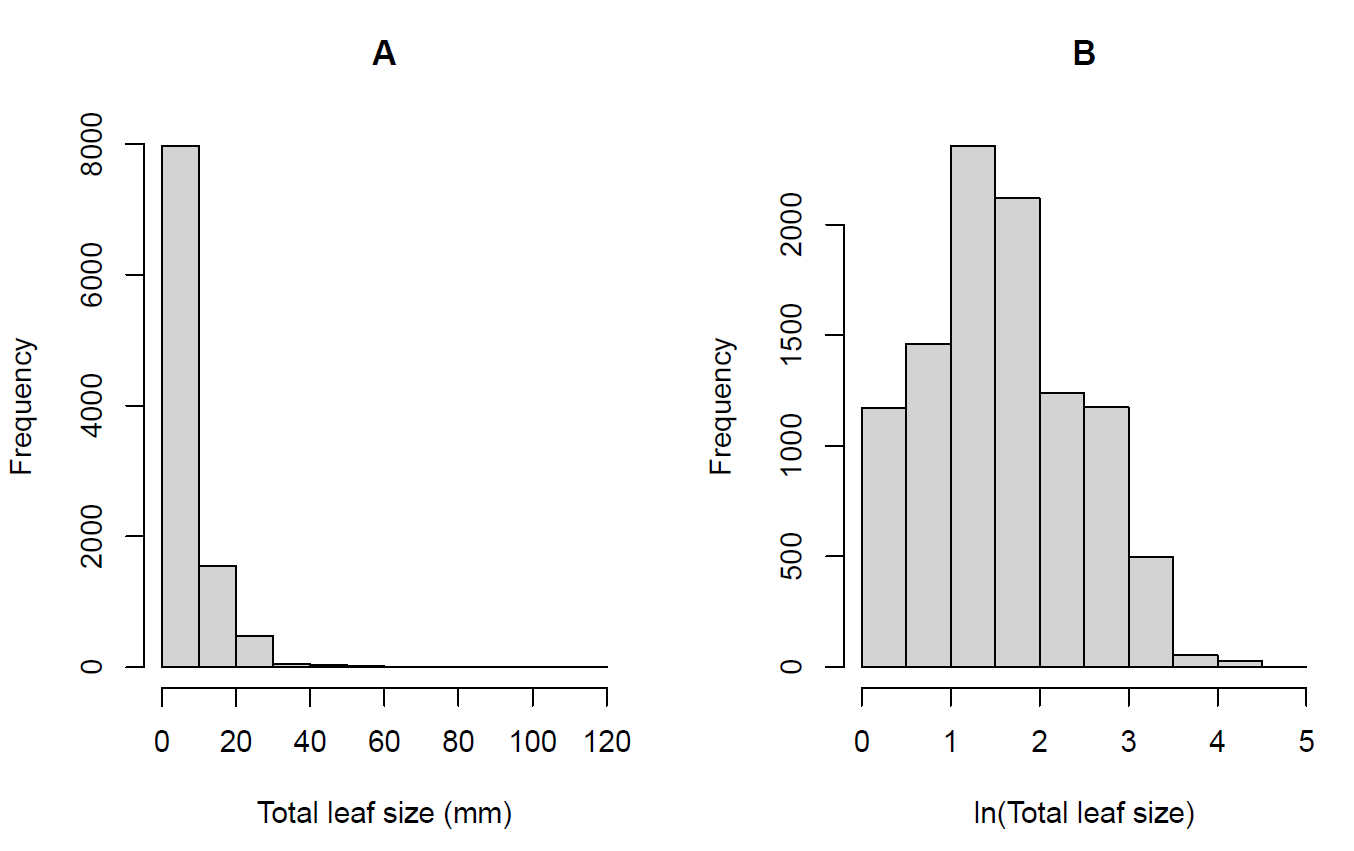

**Figure S5: Distribution of leaf sizes in the demographic dataset.** (A) Raw leaf sizes exhibited a strong right skew. (B) ln-transformed leaf sizes more closely approximated a normal distribution.
